# Telomerase reactivation induces progression of mouse Braf^V600E^-driven thyroid cancers without telomere lengthening

**DOI:** 10.1101/2023.01.24.525280

**Authors:** Iñigo Landa, Caitlin EM Thornton, Bin Xu, Jacob Haase, Gnana P. Krishnamoorthy, Jingzhu Hao, Jeffrey A Knauf, Zachary T Herbert, María A Blasco, Ronald Ghossein, James A Fagin

## Abstract

Mutations in the promoter of the telomerase reverse transcriptase (*TERT*) gene are the paradigm of a cross-cancer alteration in a non-coding region. *TERT* promoter mutations (TPMs) are biomarkers of poor prognosis in several tumors, including thyroid cancers. TPMs enhance *TERT* transcription, which is otherwise silenced in adult tissues, thus reactivating a *bona fide* oncoprotein. To study *TERT* deregulation and its downstream consequences, we generated a *Tert* mutant promoter mouse model via CRISPR/Cas9 engineering of the murine equivalent *locus* (Tert^-123C>T^) and crossed it with thyroid-specific Braf^V600E^-mutant mice. We also employed an alternative model of *Tert* overexpression (K5-Tert). Whereas all Braf^V600E^ animals developed well-differentiated papillary thyroid tumors, 29% and 36% of Braf^V600E^+Tert^-123C>T^ and Braf^V600E^+K5-Tert mice progressed to poorly differentiated thyroid cancers at week 20, respectively. Braf+Tert tumors showed increased mitosis and necrosis in areas of solid growth, and older animals from these cohorts displayed anaplastic-like features, i.e., spindle cells and macrophage infiltration. Murine *Tert* promoter mutation increased *Tert* transcription *in vitro* and *in vivo*, but temporal and intra-tumoral heterogeneity was observed. RNA-sequencing of thyroid tumor cells showed that processes other than the canonical Tert-mediated telomere maintenance role operate in these specimens. Pathway analysis showed that MAPK and PI3K/AKT signaling, as well as processes not previously associated with this tumor etiology, involving cytokine and chemokine signaling, were overactivated. Braf+Tert animals remained responsive to MAPK pathway inhibitors. These models constitute useful pre-clinical tools to understand the cell-autonomous and microenvironment-related consequences of Tert-mediated progression in advanced thyroid cancers and other aggressive tumors carrying TPMs.

## INTRODUCTION

Hotspot mutations in the proximal promoter of the telomerase reverse transcriptase gene (*TERT*) are a prototype of a non-coding genetic alteration present in multiple cancers. After the initial discovery of *TERT* promoter mutations (TPMs) in melanomas (1,2), they were also identified as frequent events in various tumor types, such as gliomas, hepatocellular, urothelial and thyroid carcinomas (3-5). Early pan-cancer studies assessing the non-coding genome identified *TERT* as the top altered gene (6,7). The former was subsequently confirmed by the Pan Cancer Analysis of Whole Genomes (PCAWG), which reported the *TERT* promoter as the most frequently mutated non-coding driver in 2,658 cancer genomes (8). Of note, TPMs correlate with metastatic and aggressive forms in most cancer lineages and have thus emerged as biomarkers of poor prognosis (1,3,9,10).

TPMs occur in two hotspots at c.-124C>T and c.-146C>T and are mutually exclusive, suggesting a common effect. Tumors harboring TPMs reactivate *TERT* transcription, which is otherwise repressed in adult normal cells. The discovery of TPMs has restored interest in TERT as a *bona fide* potential cancer target. TERT is the catalytic component of the telomerase complex and is responsible of extending chromosome ends (i.e., telomeres) by adding hexanucleotide tandem repeats, thus preventing telomere erosion and avoiding replicative senescence. In recent years, there is growing evidence of extra-telomeric roles of TERT in cancer cells, suggesting that it enhances other pro-neoplastic features (11,12).

Thyroid cancers are genetically simple tumors, typically driven by oncogenic mutations in BRAF^V600E^, *RAS* genes or receptor tyrosine kinase fusions, which result in the constitutive activation of the mitogen-activated protein kinase (MAPK) pathway. The presence of *BRAF* or *RAS* mutations in well-differentiated tumors, i.e., papillary thyroid cancers (PTC), as well as in advanced forms, such as poorly differentiated (PDTC) and anaplastic thyroid cancers (ATC), strongly suggests a continuum in disease progression via the accumulation of key additional genetic defects. Indeed, we and others reported a stepwise increase in TPM frequency along the spectrum of thyroid cancer progression: 9% in PTCs, 40% in PDTCs and 73% in ATCs. Interestingly, TPMs are subclonal in the few PTCs that harbor them, whereas they are clonal in PDTCs and ATCs, pointing to selection during tumor evolution (13-16).

So far, TERT biology in cancer has been studied in either transgenic mouse models, which do not recapitulate the endogenous levels of *Tert* expression, or in immortalized cell systems, which do not represent biologically accurate settings for telomerase biology. Early attempts to assess telomerase reactivation *in vivo* targeting Tert overexpression to various tissues showed an increase incidence of neoplastic transformation without evidence of critical telomere deregulation (17-19). Of note, Tert overexpression under the control of the keratin 5 promoter (K5-Tert) increased the frequency of pre-neoplastic lesions of the thyroid gland of aging mice, suggesting that the thyroid follicular cell lineage, even in the absence of MAPK constitutive signaling, might be particularly susceptible to telomerase reactivation (20).

In this study, we used CRISPR/Cas9 to generate a *Tert* mutant promoter mouse model and studied these animals in the context of thyroid-specific Braf^V600E^ activation. We demonstrate that Braf^V600E^+Tert^-123C>T^ mice reactivate *Tert* expression to levels comparable with those observed in human tumors harboring TPMs (e.g., BRAF^V600E^+TERT^-124C>T^), and that they induce thyroid cancer progression, mimicking the phenotypes observed in a model in which *Tert* overexpression was targeted to epithelial cells (Braf^V600E^+K5-Tert). Thyroid cells with an engineered TPM did not uniformly re-express telomerase, suggesting that additional steps are required for the enhanced *Tert* expression to manifest. Consistent with the fact that mice have longer telomeres than humans and show less restricted expression of telomerase in adult tissues (21,22), our data indicate that events unrelated to telomere attrition operate in these tumors. Instead, transcriptomic analysis of telomerase-reactivated tumors showed that overactivation of MAPK and PI3K (phosphatidylinositol-3 kinase) pathways, as well as immune-related signaling, probably in response to changes in the tumor microenvironment, likely play a role in disease progression.

## METHODS

### Sequence analysis

To determine the degree of mouse:human interspecies conservation in the region surrounding the human *TERT* c.-124C>T and c.-146C>T mutation hotspots, we performed sequence alignment between human *TERT* (chr5: 1,295,347-1,294,746) and mouse *Tert* (chr13: 73,626,701-73,627,273) covering sequences around the transcription start site using the European Bioinformatics Institute (EMBL-EBI) pairwise sequence alignment tools (https://www.ebi.ac.uk/Tools/psa/). Sequence alignments were retrieved and manually inspected for similarities.

### Luciferase assays

The 226-nucleotide sequence upstream of mouse *Tert* gene, containing the -123C wildtype allele, was synthesized at Genscript (Piscataway, NJ, USA) and cloned into the pGL4.20[luc2/Puro] reporter vector (Promega, Madison, WI, USA). A mutant version of this plasmid, containing the *Tert* -123T allele, was created by site-directed mutagenesis. *Tert* c.-123C and *Tert* c.-123T alleles were confirmed by direct sequencing. Reporter experiments were performed using the Dual-Luciferase Reporter Assays (Promega, Madison, WI, USA) following the manufacturer’s protocol and using Renilla (pGL4.73[hRluc/SV40]) as a control. *Tert* promoter-driven luciferase expression was tested in NIH-3T3 (mouse fibroblasts) and in two cell lines (B92 and B16509E) derived from murine Braf^V600E^-mutant thyroid tumors. Cells were grown in DMEM (NIH-3T3) or F12-Coon’s media (B92 and B16509E) media supplemented with 10% fetal bovine serum. Luciferase results are expressed in relative units and represent the average of three independent experiments with each condition ran in quadruplicate.

### Gene editing and Tert mutant promoter mouse generation

Two guide RNAs (gRNAs) targeting the mouse *Tert* c.-123C locus were designed. Two rounds of Cas9/CRISPR injections using the selected gRNA and donor templates were performed on mouse zygotes, followed by embryo transfer, which resulted in 84 pups. The B6 hybrid (B6CBAF1) background was used. Gene editing was performed at the MSKCC Mouse Genetics Core Facility. Animals were first screened by a restriction fragment length polymorphism (RFLP) approach, using the *Mnl*1 enzyme. Agarose gel band patterns compatible with the presence of Tert c.-123C>T mutation were amplified and Sanger-sequenced using the following primers: Tert_123_2F: 5’-CATGCACCAGCATTGTGACCA-3’ and Tert_123_4R: 5’-CAACGAGGAGCGCGGGTCATTGT-3’. To evaluate off-target effects, six founder animals with confirmed Tert -123C>T mutation were subjected to targeted next-generation sequencing (NGS) and analyzed using CRISPResso (23). NGS showed founders with up to 20% of reads displaying the desired Tert c.-123C>T mutation and no off-target effects in *cis*. Three animals were used to generate the Tert mutant promoter mouse line. Off-target alterations were bred-out in subsequent generations, as assessed by Sanger sequencing. Animals were born at the expected Mendelian frequency. The offspring of a selected founder was used to establish the Tert^-123C>T^ mouse line.

### Mouse models and breeding strategies

Animal care and all experimental procedures were approved by the MSKCC and BWH Animal Care and Use Committees. The Tert^-123C>T^ mouse line was studied in the context of thyroid-specific activation of Braf^V600E^ oncoprotein by crossing these animals with LSL-BrafV600E/TPO-Cre/eYFP mice (“Braf^V600E^”), which express endogenous levels of Braf oncoprotein in thyroid follicular cells at E14.5, when Cre recombinase is expressed downstream of the thyroid peroxidase (*Tpo*) gene promoter (24). This Braf^V600E^ model is also engineered to express yellow fluorescent protein (Jackson Lab stock #007903) in thyroid cells, as previously described (25). The transgenic Tg-K5-Tert mouse line, in which Tert expression is targeted to epithelial tissue via the keratin 5 (K5) promoter (17,20), was a generous gift from Dr. María Blasco at the Spanish National Cancer Research Centre. We crossed K5-Tert animals with our Braf^V600E^ mice to generate an alternative model in which expression of Braf^V600E^ oncoprotein and overexpression of Tert were present in thyrocytes.

Our breeding scheme was designed to minimize housing differences between the groups to be compared, ensuring that Braf^V600E^ animals, and either Braf^V600E^+ Tert^-123C>T^ or Braf^V600E^+K5-Tert mice were littermates. Tert c.-123C>T was studied in heterozygosis. Braf^V600E^ alone was used as our baseline model of well-differentiated thyroid cancer for all comparisons. Mouse genotyping was primarily achieved by allele-specific assays designed and performed by Transnetyx.

### Histology and Immunohistochemistry

Thyroid tissues were fixed in 4% paraformaldehyde, embedded in paraffin, sectioned, and stained with hematoxylin and eosin (H&E). Histological diagnosis was performed by thyroid pathologists (R.G. and B.X.) blinded to the genotype and the treatment status of each animal. Immunohistochemistry (IHC) was performed on the Leica Bond III automated staining platform using the Leica Biosystems Refine Detection Kit. The following antibodies were used for IHC: phospho-Erk (Cell Signaling Technology (CST), cat. #4370), Ki-67 (CST, cat. #12202), CD11b (Abcam, cat. #ab133357), F4/80 (CST, cat. #70076), Arg1 (Abcam, cat. #ab 93668) and Pax8 (Proteintech, cat. #10336). The secondary antibodies were part of the Leica Bond Polymer Refine Detection Kit (Catalog No: DS9800). QuPath (Version 0.3.0, (26)) was used for H/E slide visualization and the quantification of DAB (3,3⍰-Diaminobenzidine) staining on IHC slides. Tumor boundaries were delineated, and the positive cell detection command was applied to the tumor area, applying default settings and a single intensity threshold of 0.2 to quantify the percentage of positive DAB-stained cells.

### Generation of mouse thyroid cancer cell lines

To generate mouse cancer cell lines, thyroid tumors were dissected from surrounding tissues and collected on minimum essential media (MEM). Specimens were subsequently minced using sterile razorblades, spun down by centrifugation at 500 x g for 5 minutes and resuspended in 10 ml of digestion medium (MEM containing 112 U/mL type I collagenase; Worthington, cat. #CLS-1), 1.2 U/mL dispase (Gibco; catalog no. 17105-041), penicillin (50 U/mL), and streptomycin (50 mg/mL). Cells were incubated at 37°C for 60 minutes with vigorous shaking, after which cells were spun down and resuspended in Coon’s modified F12 medium with penicillin/streptomycin/L-glutamine (P/S/G; Gemini; #400-110) and 0.5% bovine brain extract (BBE, Hammond Cell Tech, cat. #2007-NZ), plated into CellBind plates (Corning Inc.) for two weeks and then switched to Coon’s modified F12 medium with P/S/G containing 5% FBS for routine culturing and subsequent experiments. Cell lines were maintained at 37°C and 5% CO2 in humidified atmosphere.

### Flow cytometry

To isolate YFP+ thyroid cells for subsequent RNA sequencing (RNAseq), tumor and normal thyroids were harvested in cold digestion buffer (Hank’s Balanced Salt Solution (HBSS) supplemented with 5% FBS and 1.5 mg/ml Collagenase A, and thoroughly minced with razorblades. Tissue suspensions were transferred to 15 ml tubes and incubated at 37°C for 1 hour with intermittent vortexing every 10 minutes. Dissociated thyroid cells were passed through a 70 µm cell strainer, pelleted by centrifugation at 500 × g for 5 minutes, washed twice with cold PBS and resuspended in cold sorting buffer (F-12 Coon media containing 2% FBS). A YFP+ pure thyroid cell population was sorted using a BD FACSAria flow cytometer into TRIzol (Invitrogen, #15596018). Each mouse tumor from Braf^V600E^, Braf^V600E^+Tert^-123C>T^ and Braf^V600E^+K5-Tert mice, was processed individually, and target cell yield was established at 20,000 YFP+ cells. For non-Braf specimens which did not generate tumors, we pooled several normal-looking thyroid glands from identical genotypes as follows: wildtype (WT, n=25), Tert^-123C>T^ only (n=17) and K5-Tert only (n=14).

### RNA sequencing

Total RNA was isolated from up to 20,000 YFP+ cells sorted into TRIzol. Phase separation in cells lysed in TRIzol reagent was achieved with chloroform. RNA was precipitated with isopropanol, washed with 75% ethanol, and resuspended in RNase-free water following the manufacturer’s protocol. RNAseq was performed at the Dana-Farber Cancer Institute Molecular Biology Core Facility. RNA libraries were prepared following a low input mRNAseq protocol. RNA samples were fragmented at 94°C for 8 min with 14 cycles of PCR post-adapter ligation, according to manufacturer’s recommendation. The finished dsDNA libraries were quantified by Qubit fluorometer and Agilent TapeStation 2200. Libraries were pooled in equimolar ratios. Final sequencing was performed on an Illumina NovaSeq with paired-end 100 bp reads.

Standard analysis and visualization of RNAseq data was performed using VIPER (27). These include the generation of heatmaps and volcano plots, as well as pathway and functional group analysis employing Gene Ontology (GO), Gene Set Enrichment Analysis (GSEA) and the Kyoto Encyclopedia of Genes and Genomes (KEGG) pathway database. Expression of specific genes was evaluated from RNAseq-derived normalized counts and represented as median ± interquartile range (IQR) of gene transcripts for each animal. Specific pairwise comparisons from RNAseq data were performed by comparing medians of normalized counts from defined groups, as described, and *P* values were calculated using unpaired two-tailed Mann-Whitney U tests, unless otherwise noted.

### Quantitative real-time PCR

RNA was preserved in TRIzol (Invitrogen) and isolated as described above. One microgram of total RNA per sample was reverse-transcribed into cDNA using SuperScript III Reverse Transcriptase (Invitrogen). Quantitative, real-time PCR (qPCR) was carried out in triplicates using Power SYBR Green PCR Master Mix (Applied Biosystems). Gene-specific, intron-spanning primer pairs for qPCR were designed for each target, and β-actin gene was used as housekeeping control for data normalization and qPCR analysis, using the delta-delta Ct method. Primer list is available in Supplementary Table S1.

### Western Blotting

Thyroid lobes were surgically resected under the microscope to ensure the removal of surrounding tissues and immediately placed in liquid nitrogen for preservation until protein extraction. Cultured cells were harvested with 0.05% trypsin/0.02% EDTA solution, and cell pellets were washed with cold PBS. Proteins were extracted using RIPA buffer (EMD Millipore) supplemented with protease and phosphatase inhibitors as per the manufacturer’s instructions. Protein concentrations were estimated by the BCA Kit (Thermo Scientific) on a microplate reader (SpectraMax M5). Comparable amounts of proteins were subjected to SDS-PAGE using NuPAGE 4% to 12% Bis–Tris gradient gels (Invitrogen) and were transferred to PVDF membranes. Following overnight incubation with the primary antibody at 4°C, membranes were incubated with goat anti-rabbit or goat anti-mouse secondary antibodies coupled to horseradish peroxidase (HRP) for 1 hour at room temperature. Chemoluminescence was detected using Pierce ECL Western Blotting Substrate (Thermofisher Scientific) and visualized on a ChemiDoc equipment. The following primary antibodies were used: phospho-Erk (CST, cat. #4370), total Erk (CST, cat. #4695), phospho-Akt (at Ser473, CST, cat. #4060), total Akt (CST, cat. #4691), PI3 Kinase p85 (CST, cat. #4292), phospho-NFκB p65 (at Ser536, CST cat. #3033) and vinculin (CST cat. #13901). Quantification of western blot bands was done using ImageJ Fiji (28) and normalizing phospho-proteins to loading controls.

### In situ hybridization of single RNA molecules by RNAscope

In situ identification of messenger RNA (mRNA) molecules at single-cell resolution on mouse thyroid specimens was performed in conjunction with Harvard Medical School Neurobiology Imaging Facility using RNAscope. We evaluated a total of 20 murine thyroid specimens harvested from 20-week animals representing the following genotypes: wildtype (n=2), Braf^V600E^ (n=5), Braf^V600E^+Tert^-123C>T^ (n=8) and Braf^V600E^+K5-Tert (n=5). These thyroid specimens were FFPE-embedded and mounted on slides, ensuring that different genotypes were included on the same slide, and subsequently processed for RNAscope. We followed the standard protocol for RNAscope multiplex fluorescent reagent kit v2 assay (cat. #323100, ACD-BioTechne). Tert mRNA was detected using RNAscope fluorescent probe cat. #313441. Images were acquired on an Olympus VS120 Whole Slide Scanner with Hamamatsu Orca R4 camera and 40X 0.95NA objective. RNAscope image analysis was performed using QuPath V 0.3.2 (26). Image files in

*.ets format were uploaded using the Bio-Formats builder. Tumor boundaries were outlined manually, and individual cells were detected via DAPI staining and counted using the automated cell detection algorithm. Subcellular detection was performed to identify RNA transcripts in the Cy5 or TRITC channels. The thresholds <1000 and <8000 were input for Cy5 and TRITC, respectively, and the size parameters of 4.5 µm2 average area per spot was input. Quantification of Tert mRNAs across specimens was performed by a single operator (CEMT) who was blinded to the tumor genotypes. Qupath-generated data on the number of RNAscope spots per cell was exported into Excel and genotypes were only unblinded for comparison between genotypes. Each specimen was mounted in duplicate on each slide and percentages of cells expressing Tert transcripts were averaged. Data was presented as the percentage of all cells within each specimen expressing 1, 2, 3, 4 and 5+ (range 5-10) fluorescent spots, representing single-cell level Tert mRNAs.

### Thyroid ultrasonic imaging

Mice were anesthetized by inhalation of isoflurane. Thyroid tumors were imaged using Vevo 770 High-Resolution *In Vivo* Micro-Imaging System (VisualSonics). Aqueous ultrasonic gel was applied to the denuded skin overlying the thyroid gland prior to placement of the ultrasonic transducer. Volume was calculated by manually tracing the margin of the tumor every 250 μm using the instrument software. Genotypes were blinded for volume analysis. Volume calculation for all tumors was performed by the same person (IL) to avoid inter-operator bias.

### Drug administration in vivo

A cohort of 24 animals, including eight Braf^V600E^, eight Braf^V600E^+Tert^-123C>T^ and eight Braf^V600E^+K5-Tert mice were aged for 20 weeks and randomized into two balanced groups within each genotype: “vehicle” and “treated” (n=4 per genotype and condition). The treated group was administered a combination of the Raf inhibitor dabrafenib (GSK2118436, Selleckchem) at 30 mg/kg plus the Mek inhibitor trametinib (GSK1120212, Selleckchem) at 3 mg/kg. Drugs were prepared in 0.5% HPMC, 0.2% Tween80 in H2O pH 8.0, and administered daily by oral gavage. The vehicle cohort was given the drug solvent by oral gavage under the same conditions. Animals were treated daily for 12 days, with the weekend off drugs. Thyroid volumes were monitored by ultrasound on days 1 and 12. Mouse weight was recorded on days 2, 5, 8, 10 and 12. All animals were sacrificed on day 12, within two hours from the last dose. Thyroid glands were collected under the microscope to ensure the removal of surrounding tissues. One lobe was fixed and paraffin-embedded for histological and IHC analyses. The other lobe was minced and preserved in TRIzol for RNA extraction.

### Telomere length measurement

Telomere length from mouse thyroid primary tumors and tumor-derived cell lines was measured as previously described (29). Briefly, thyroid tumors from each genotype from 20-week-old animals were flash-frozen. Frozen tumors were mechanically homogenized in DNeasy (Qiagen) extraction kit buffer, and genomic DNA (gDNA) was isolated following the manufacturer instructions. For cell line experiments, cells derived from age-matched animals were grown until a similar passage, and gDNA was isolated as explained above. Up to 30 nanograms of gDNAs were used in equal amounts for quantitative PCR-based measurement of telomeres, using primers for mouse telomeric region and *36b4* as the housekeeping gene, as reported (29). Reactions were run in quadruplicate wells, and results were normalized across two independent experiments.

### Statistical analysis and data graphing

GraphPad Prism software (version 9.3.1) was used for statistical analyses and graphics generation. Data from qPCR, ultrasonic imaging, quantification of immunohistochemistry and RNAscope images are represented as either mean ± standard deviation (SD), or median ± interquartile range (IQR), depending on whether distributions passed normality tests. Consequently, *P* values for pairwise comparisons were calculated using either unpaired two-tailed student t tests or Mann-Whitney U tests, as noted. Three-group comparisons were performed by either ANOVA or Kruskal-Wallis tests. Conceptual figures describing experimental approaches were created with BioRender.com.

### Data availability statement

The RNA-seq data generated in this study are publicly available in Gene Expression Omnibus (GEO) at GSEXXXXX.

## RESULTS

### Mutation at mouse *Tert* c.-123C>T is conserved and mimics the transcriptional effects of human *TERT* c.-124C>T

We first checked whether the *TERT* proximal promoter region, where hotspot mutations occur in human tumors, was conserved in the mouse. Pairwise sequence alignments showed that the human *TERT* site around -124C>T mutation, which accounts for ∼80% of TPMs in human tumors (3), was conserved in murine sequences. We identified a mouse promoter residue at -123C (from the ATG codon), whose flanking nucleotides align well with the human ortholog, and which upon a C>T substitution create an identical core binding sequence for ETS factors, as defined in both species (30) (Fig 1A, Suppl Fig S1). We then tested if a mouse *Tert* promoter construct harboring the -123C>T mutation led to increased *Tert* expression in luciferase assays performed in NIH-3T3 (mouse fibroblasts) and two cell lines derived from murine Braf ^V600E^-mutant thyroid tumors. All three mouse cell lines showed a two to three-fold increase in *Tert* promoter activity for the mutant promoter (“mTert -123T”) compared to wildtype (“mTert - 123C”) constructs (Fig 1B). This is consistent with the effect seen in luciferase assays for human *TERT* - 124C>T/-146C>T constructs (31), as well as with TERT expression in patients’ tumors carrying TPMs (32,33).

**Figure 1.**
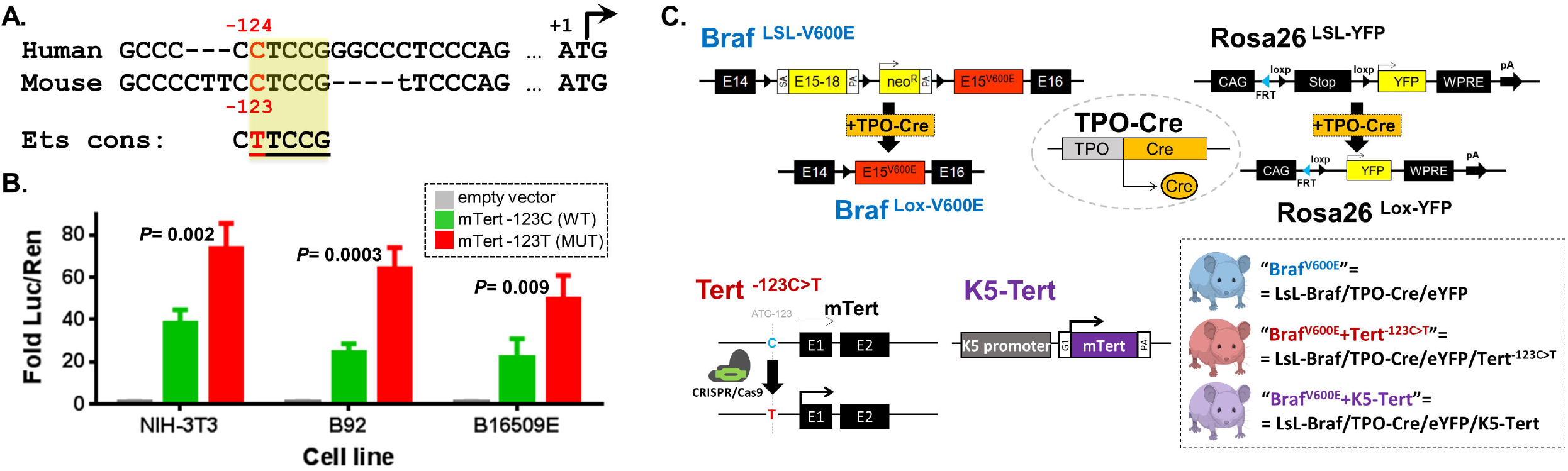
Modeling of Tert reactivation in genetically engineered mouse models with endogenous Braf^V600E^ expression. **A**. Schematic representation of the sequence alignment of human *TERT* (top) and mouse *Tert* promoter (bottom) sequences. A single C>T transition at the conserved human c.-124C and mouse c.-123C *loci* generates a consensus binding motif for Ets transcription factors (“Ets cons”); *B*. Tert promoter-driven luciferase expression, normalized by renilla (“Luc/Ren”), in mouse cell lines using Tert wildtype (mTert -123C, green) and mutant (mTert -123T, red) constructs; **C**. Genetic schema of the mouse models: Top: Thyroid-specific expression of Cre recombinase (driven by thyroid peroxidase (TPO) in TPO-Cre constructs) substitutes exon 15 of WT Braf by V600E mutant allele, resulting in endogenous expression of Braf oncoprotein (66). Cre-mediated excision of stop cassette also enables YFP (“yellow fluorescent protein”) expression in thyroid cells; Bottom: Tert^-123C>T^ is knocked-in in the germline via CRISPR/Cas9 editing of mouse zygotes, as described (left); mouse Tert is driven by the keratin 5 (K5) promoter (middle) (17); breeding of LsL-Braf, TPO-Cre/eYFP and Tert alleles generates the three genotype combinations used in this study (right).

### Generation of a Tert mutant promoter mouse model

To investigate the role of *Tert* promoter mutations *in vivo*, we generated a Tert c.-123C>T mouse model by Cas9/CRISPR gene editing (“Tert^-123C>T^” from now on). We successfully targeted the region in mouse zygotes and obtained 10/84 F0 animals carrying the desired -123C>T mutation. Targeted NGS around the *Tert* promoter locus on six F0 animals confirmed the knock-in of Tert c.-123C>T mutation without off-target effects in *cis* in 1-20% of the sequencing reads (Suppl Table S2). Off-target mutations typically consisted of single nucleotide changes or indels around the desired residue and were bred-out in the subsequent generations. A mouse Tert c.-123C>T line without any off-target alterations was established and confirmed by direct sequencing of the offspring. We subsequently crossed Tert^-123C>T^ animals with LSL-BrafV600E/TPO-Cre/eYFP mice (“Braf^V600E^”), which express endogenous levels of the Braf oncoprotein and yellow fluorescent protein (YFP) in thyroid follicular cells. In parallel, we crossed mice in which Tert expression was targeted to epithelial tissues via the keratin 5 promoter (“K5-Tert”) with the same thyroid-specific Braf^V600E^ model (Fig. 1C). All animals were viable and born at the expected Mendelian rates >

### Mice carrying Braf ^V600E +Tert -123C>T^ alleles develop advanced thyroid cancers

We monitored tumor progression in Braf^V600E^ *vs*. Braf^V600E^+Tert^-123C>T^ animals and employed the Braf^V600E^+K5-Tert mice as positive controls of sustained activation of Braf and Tert oncoproteins in thyroid follicular cells. Braf^V600E^ mice developed PTCs at 4-6 weeks with nearly 100% penetrance, as previously reported (24), but they hardly ever progressed to advanced disease. We first sacrificed a cohort of ∼10-week-old Braf^V600E^ and Braf^V600E^+Tert^-123C>T^ animals (Figure 2A). All Braf-mutant animals developed PTCs, characterized by the presence of papillae and specific nuclear features (Suppl Fig S2A), and so did most of the Braf^V600E^+Tert^-123C>T^ mice, except 1/12 Braf^V600E^+Tert^-123C>T^ animals, which displayed a PDTC phenotype (Table 1). We then monitored a larger cohort of mice for an extended period and confirmed that a subset of Braf^V600E^+Tert^-123C>T^ animals progressed to histologically more aggressive tumors at around 20 weeks. As shown in Table 1 and Figure 2B-C, none of the Braf^V600E^ animals in this age range developed PDTCs (0/12), whereas 7/24 (29.2%) Braf^V600E^+Tert^-123C>T^ mice did (chi-squared p-value= 0.0371). Braf^V600E^+Tert^-123C>T^ tumors typically preserved a PTC component and displayed areas of PDTC (Figure 2B). A subset of these Braf^V600E^+Tert^-123C>T^ tumors showed characteristics compatible with the Turin definition of PDTC (Figure 2B, second panel), whereas most fulfilled the criteria for the “high-grade differentiated thyroid carcinoma” (HGDTC) category (Figure 2B, third panel) (34-36). Interestingly, a similar phenotype was observed in an age-matched cohort of the Tert overexpression transgenic model (Braf^V600E^+K5-Tert): 4 out of 11 (36.4%) developed PDTCs (Figure 2B, bottom panel and Figure 2C). We noted other histological features in these tumors. Muscle invasion was observed in 25.0, 37.5 and 45.5% of Braf^V600E^, Braf^V600E^+Tert^-123C>T^ and Braf^V600E^+K5-Tert animals, respectively. In Braf+Tert tumors, areas of solid growth correlated with proliferating cells and the presence of mitotic figures (Suppl Fig 2B). Braf+Tert animals tended to have larger tumors and diminished survival, but differences did not reach statistical significance (Suppl Fig S2C-D). The presence of inflammation and calcifications were observed only in 20-week Braf^V600E^+Tert^-123C>T^ and Braf^V600E^+K5-Tert groups.

**Table 1.** Histological features of mouse thyroid tumors.

| Genotype | 6-12 weeks |  |  |  |  | 20-30 weeks |  |  |  |  |
| --- | --- | --- | --- | --- | --- | --- | --- | --- | --- | --- |
|  | Age - avg weeks | PTC - n (%) | PDTC - n (%) | p-value | Total - n | Age - avg weeks | PTC - n (%) | PDTC - n (%) | p-value | Total - n |
| **Braf<sup>V600E</sup>** | 9.9 | 8 (100.0) | 0 (0.0) | 0.4215 | 8 | 22.2 | 12 (100.0) | 0 (0.0) | - | 12 |
| **Braf<sup>V600E</sup> +Tert<sup>123C>T</sup>** | 10.2 | 12 (93.3) | 1 (7.7) |  | 13 | 22.0 | 17 (70.8) | 7 (29.2) | 0.0371 | 24 |
| **Braf<sup>V600E</sup> +K5-Tert** | N/A | N/A | N/A | N/A | N/A | 23.7 | 7 (63.6) | 4 (36.4) | 0.0765 | 11 |

Abbreviations: PTC=papillary thyroid cancer; PDTC=poorly differentiated thyroid cancer; p-values are derived from 2-sided chi-square tests for the indicated groups, employing Braf<sup>V600E</sup> animals as controls.

**Figure 2.**
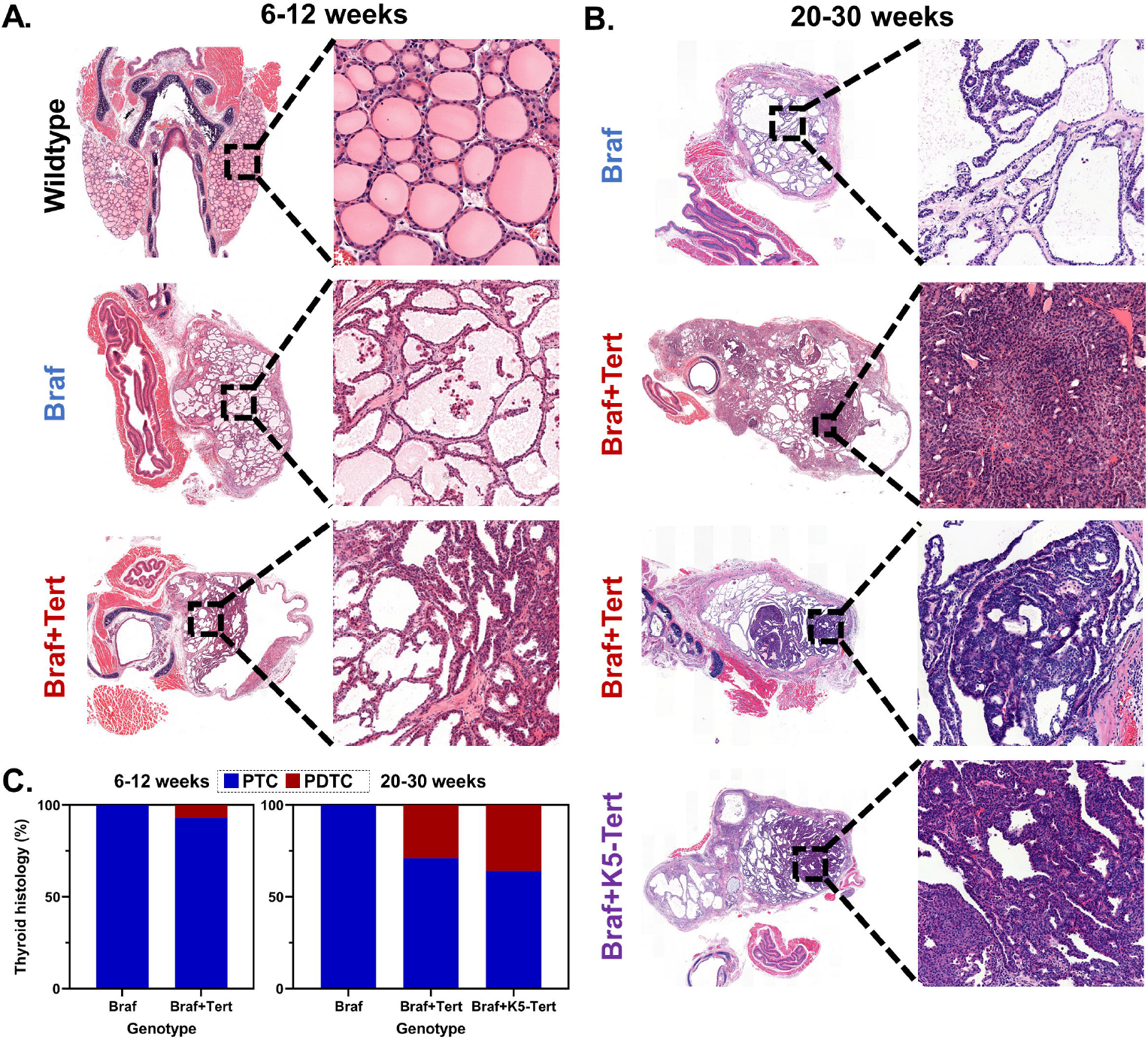
Histological analysis of the Braf+Tert mouse models. Representative H&E-stained thyroid sections of Braf^V600E^, Braf^V600E^+Tert^-123C>T^ and Braf^V600E^+K5-Tert mice at **A**. 6-12 weeks; and **B**. 20-30 weeks. **C**. Percentage of animals with the indicated genotypes showing papillary (PTC) or poorly differentiated thyroid cancers (PDTC) at 6-12 weeks (left) and 20-30 weeks (right). Abbreviations: “Braf+Tert” and “Braf+K5-Tert” labels denote “Braf^V600E^+Tert^-123C>T^” and “Braf^V600E^+keratin 5-driven Tert^”^ genotypes, respectively.

We then followed a small cohort of Braf^V600E^ and Braf^V600E^+Tert^-123C>T^ animals up to 40 weeks. At that time point, a subset of mice from both groups showed more advanced phenotypes; however, only Braf^V600E^+Tert^-123C>T^ mice displayed ATC-like features, including pleomorphic or spindle nuclei, loss of positivity for Pax8 and macrophage infiltration (Suppl Fig S2E). These same features were also observed in some 20-week-old Braf+K5-Tert mice, suggesting that high level expression of Tert might cooperate with oncogenic Braf to promote PTC-to-ATC transformation (Suppl Fig S2F). Tumor infiltration by myeloid cells was observed in 40-week-old Braf^V600E^ and Braf^V600E^+Tert^-123C>T^ animals; however, a subset of Braf^V600E^+Tert^-123C>T^ tumors, displayed the highest proportion of cells staining positive for markers of M2-like macrophages, thus mimicking a hallmark of human ATCs (Suppl Fig S2G-H).

Overall, Tert reactivation accelerated Braf-driven PTC progression towards more advanced forms, as observed in patients’ tumors carrying this genetic combination. Our results suggested that 20 weeks was an optimal window to evaluate Tert-mediated changes on thyroid cancer progression, so we used this time point for all subsequent experiments.

### Braf^V600E^ +Tert^-123C>T^ tumors ultimately acquire the ability to increase Tert transcription, but Tert reactivation displays temporal and intra-tumoral heterogeneity

We then tested whether the engineering of a Tert^-123C>T^ mutation led to an increase in Tert transcription *in vivo*. Tert mRNA levels of YFP-sorted cells from thyroid tumors of 10-week Braf^V600E^+Tert^-123C>T^ animals were similar from those from Braf^V600E^ mice, which, unlike human tissues, retain baseline Tert expression (21). However, at 20 weeks, Tert mRNA levels of a subset of Braf^V600E^+Tert^-123C>T^ increased and were higher than 10-week-old specimens with the same genotype (Fig 3A, *P*= 0.006). This suggests that Tert re-expression tracks with progression from PTC to PDTC, probably because cells with higher Tert levels are clonally selected over time. To further dissect the heterogeneity of Tert expression in animals engineered for the promoter mutation, we evaluated the levels of Tert transcripts *in situ* at single-cell resolution via RNAscope analysis of 20-week thyroid tumors from each genotype. A subset of thyroid cells from wildtype and Braf^V600E^ mice maintained some Tert expression, as expected. Not surprisingly, Braf+K5-Tert animals showed sustained, overexpression of Tert mRNAs in about half of thyroid cells (Fig 3B). Interestingly, the average number of Tert transcripts per cell was higher in Braf^V600E^+Tert^-123C>T^ tumors compared to Braf^V600E^ for cells displaying high Tert transcription (Fig 3C, see “3”, “4” and “5+” transcript categories). In addition, the distribution of Tert transcripts in Braf^V600E^+Tert^-123C>T^ animals showed an intra-tumoral heterogeneity which was not observed in Braf^V600^ tumors. Crucially, areas of solid growth identified as PDTC regions typically correlated with a much higher number of fluorescent Tert mRNAs (Suppl Fig S3).

**Figure 3.**
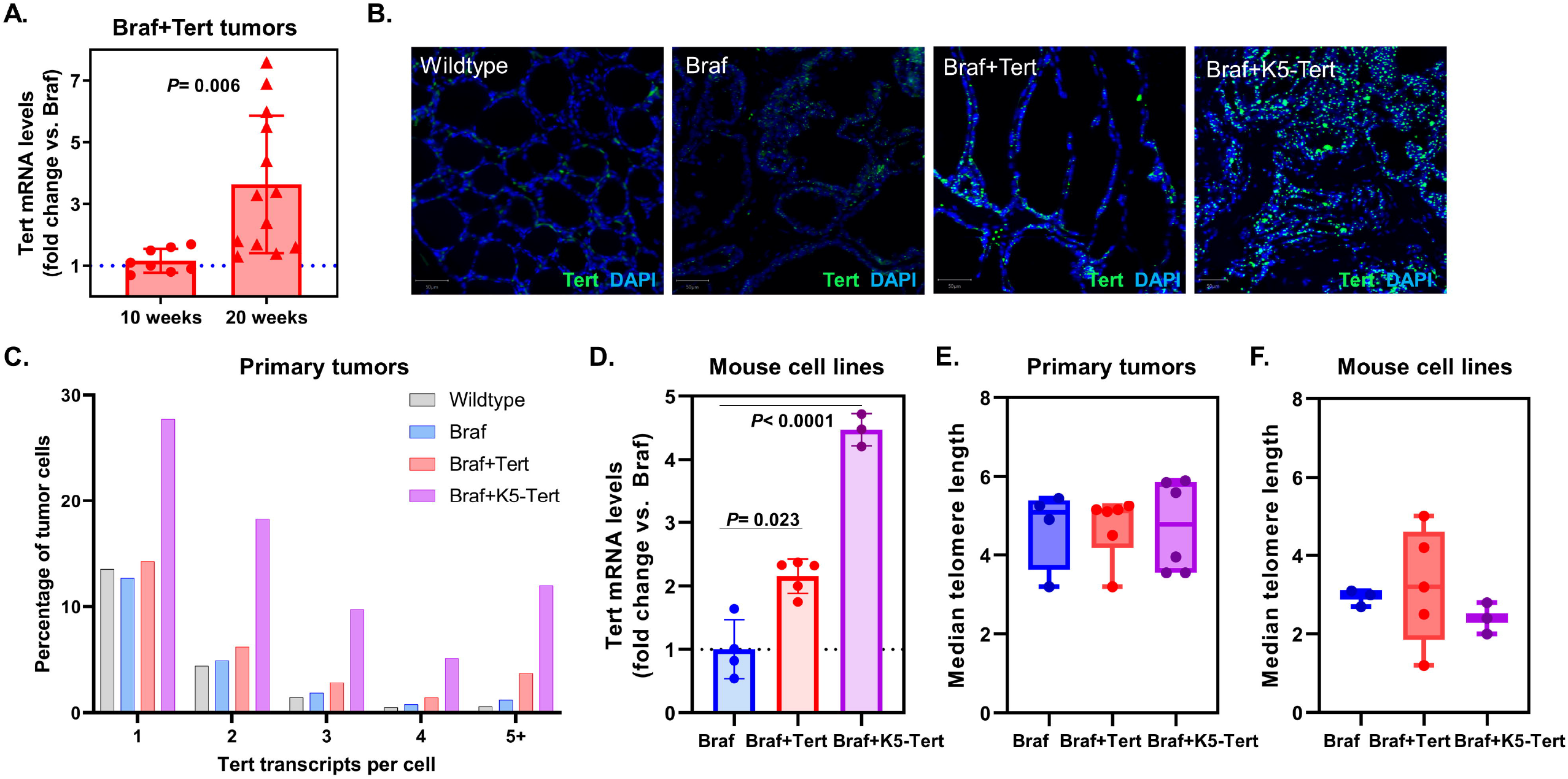
Characteristics of Tert re-expression in telomerase-reactivated thyroid cancers. **A**. Relative levels of Tert transcription in YFP-sorted cells isolated from Braf^V600E^+Tert^-123C>T^ mouse tumors, showing an increase in Tert mRNA levels for specimens collected at 20 weeks *vs*. 10 weeks. Each red point represents a tumor collected from a different animal at the indicated age. Results are expressed as fold change compared to the Tert baseline expression of Braf^V600E^ tumors from the same ages (dotted blue line). **B**. Representative examples of tumors from 20-week animals with the indicated genotypes subjected to RNAscope to detect Tert single mRNA molecules (green dots). DAPI (blue) is used for contrast. **C**. Quantification of number of Tert transcripts at single-cell resolution from RNAscope data on mouse tumors with the indicated genotypes. Data is represented as percentage of cells within each tumor expressing 1, 2, 3, 4, or 5+ transcripts. Tumor boundaries were defined manually and cell detection and quantification of green fluorescent dots was performed employing the built-in automated analysis tools on QuPath, using the same detection parameters across specimens. **D**. Relative mRNA levels in cell lines derived from Braf^V600E^ (n=4), Braf^V600E^+Tert^-123C>T^ (n=5) and from Braf^V600E^+K5-Tert (n=3) mouse tumors, showing that genotype-dependent increases in Tert transcription are maintained *in vitro*. **E**. Median telomere length, expressed in relative units, of cell lines derived from mouse thyroid tumors with the indicated genotypes. **F**. Median telomere length, expressed in relative units, of YFP-sorted cells isolated from mouse thyroid tumors with the indicated genotypes. Abbreviations: “Braf+Tert” and “Braf+K5-Tert” labels denote “Braf^V600E^+Tert^-123C>T^” and “Braf^V600E^+keratin 5-driven Tert^”^ genotypes, respectively.

We also derived primary cultures from these tumors, which grew well over multiple passages. Interestingly, these ***in vitro*** models maintained their original genotype-dependent differences in Tert expression: Braf^V600E^+Tert^-123C>T^ and Braf+K5-Tert showed a 2.1- (***P***= 0.023) and 4.5-fold (***P***<0.0001) increase in Tert transcription, respectively, ***vs***. Braf cells (Fig 3D). This points to Braf^V600E^+Tert^-123C>T^ lines, in which presumably only a subset of cells from the original specimens increased Tert mRNA levels, selecting for Tert overexpression during ***in vitro*** immortalization.

### Telomere length is not altered in tumors with Tert reactivation

We then explored whether engineered re-expression of Tert, either via c.-123C>T mutation or keratin 5 promoter-mediated transcription, affected telomere length in mouse primary tumors. Relative median ± interquartile range (IQR) telomere length in Braf^V600E^, Braf^V600E^+Tert^-123C>T^, Braf^V600E^+K5-Tert thyroid tumors was 5.08±1.78, 5.13±1.05 and 4.78±2.31, respectively (Kruskal-Wallis p-value= 0.8087; Fig 3E). This suggested that telomere attrition is likely not a critical factor in the observed murine Tert-driven thyroid cancer progression. Our findings are in line with previous reports in which telomerase overexpression induced cancers without critical changes in telomere length (18-20), as well as the known fact that mice have unusually long telomeres compared to humans (22). In addition, we evaluated telomere length in immortalized cell lines that we derived from these murine tumors, confirming the lack of differences in specimens from each genotype evaluated at a similar passage: median±IQR was 3.20±0.40, 3.20±2.75 and 2.40±0.80, in Braf^V600E^, Braf^V600E^+Tert^-123C>T^, Braf^V600E^+K5-Tert cell lines, respectively (Kruskal-Wallis p-value= 0.3929; Fig 3F). Overall, these findings prompted us to look for non-telomeric effects as the primary mechanisms driving the observed Tert-induced thyroid cancer progression in our models.

### Thyroid tumors with telomerase reactivation display distinct transcriptomes

To evaluate whether telomerase reactivation impacted the transcriptome of Braf-driven cancers, we ran RNAseq on YFP-sorted cells from primary thyroid tumors from unselected 20-week Braf^V600E^, Braf^V600E^+Tert^-123C>T^ and Braf^V600E^+K5-Tert animals (Fig 4A). For comparison purposes, we also ran RNAseq on YFP+ thyroid cells from pooled thyroid glands from WT, Tert^-123C>T^ and K5-Tert animals (without oncogenic Braf activation), which, as expected, did not develop tumors. RNAseq normalized counts for Tert were increased in pooled YFP+ cells from Tert^-123C>T^ and K5-Tert thyroids compared to WT (Fig 4B). When comparing tumor-bearing animals, Braf^V600E^ mice showed high baseline expression of Tert (compared to WT). Tert constitutive upregulation was observed in Braf^V600E^+K5-Tert (***P***= 0.029, compared with Braf^V600E^), but not in Braf^V600E^+Tert^-123C>T^ (Fig 4C), consistent with the evidence that only certain cells within the tumor engineered for Tert c.-123C>T mutation overexpress this gene (Fig 3B-C and Suppl Fig S3).

**Figure 4.**
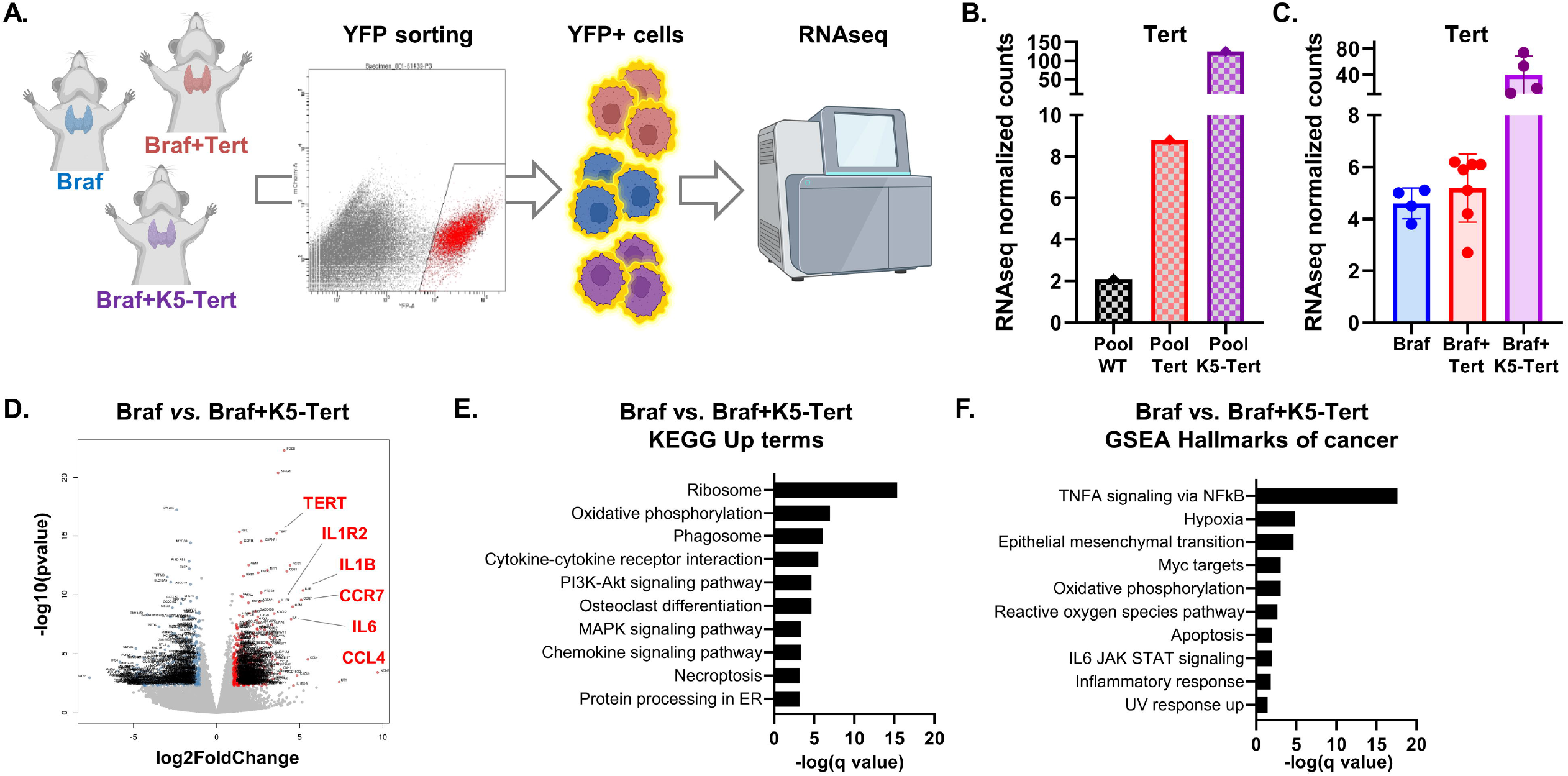
Transcriptomic characterization of telomerase-reactivated mouse thyroid tumors. **A**. Schematic representation of the isolation of thyroid tumor cells for RNA sequencing (RNAseq). **B**. Comparison of RNAseq normalized counts for Tert in pooled thyroid glands with the indicated genotypes. Values represent average expression levels for pooled WT (n=25), Tert^-123C>T^ (n=17) and K5-Tert (n=14) animals. **C**. Comparison of RNAseq normalized counts for Tert in thyroid tumors from Braf^V600E^ (n=4), Braf^V600E^+Tert^-123C>T^ (n=7) and Braf^V600E^+K5-Tert (n=4) animals. Each dot represents an individual mouse. **D**. Volcano plot showing significantly under- and overexpressed genes in Braf^V600E^+K5-Tert tumors compared to Braf^V600E^. Specific genes are indicated in red. **E**. Top 10 upregulated terms from the KEGG (Kyoto Encyclopedia of Genes and Genomes) pathway database analysis for the Braf^V600E^ vs. Braf^V600E^+K5-Tert comparison. **F**. Top 10 upregulated terms from the GSEA (Gene Set Enrichment Analysis) hallmarks of cancer analysis for the Braf^V600E^ vs. Braf^V600E^+K5-Tert comparison.

We subsequently assessed global gene expression changes in these specimens. Unsupervised clustering classified tumors primarily based on their genotypes (Suppl Fig S4A). Bulk RNA sequencing identified only a small subset of genes differentially expressed in the Braf^V600E^+Tert^-123C>T^ models compared to Braf^V600E^ (Suppl Table S3), as changes were likely diluted by the observed intratumoral heterogeneity of Tert reactivation (Fig 3B-C). The amount and extent of differences were more pronounced for Braf^V600E^+K5-Tert than for Braf^V600E^+Tert^-123C>T^, so we primarily used keratin 5-mediated Tert overexpression models to inform our subsequent analyses (Fig 4D and Suppl Table 4).

Braf^V600E^+K5-Tert tumors showed distinct transcriptomes, with multiple under- and over-expressed genes, compared to their Braf^V600E^ counterparts (Fig 4D). In addition to Tert, genes encoding several cytokines, chemokines and some of their receptors, were overexpressed in these specimens (see highlighted genes in Fig 4D). Pathway analysis confirmed these observations: cytokine and chemokine signaling were among the top overexpressed terms in Braf^V600E^+K5-Tert tumors when employing the Kyoto Encyclopedia of Genes and Genomes (KEGG) pathway database (Fig 4E and Suppl Table S5). In addition, the top two upregulated terms by Gene Ontology (GO) analysis of this same dataset were “immune system process” (q-value<1E-60) and “inflammatory response” (q-value<1E-40). The paucity of transcriptomic changes observed in Braf^V600E^+Tert^-123C>T^ by bulk RNAseq prevented us from running in-depth pathway analysis for these specimens. However, it is worth noting that the top two KEGG upregulated terms (employing unadjusted p-values) in Braf^V600E^+Tert^-123C>T^, compared to Braf^V600E^, were “chemokine signaling” and “cytokine-cytokine receptor interaction”, pointing to a similar signaling hub regardless of the mechanisms by which we induced Tert re-expression (Suppl Fig 4B and Suppl Table S6). Seeking to more precisely pin down the signaling pathways by which these cytokines might be relevant in thyroid tumors, we applied the gene set enrichment analysis (GSEA) hallmarks of cancer database analysis to our RNAseq dataset. As shown in Fig 4F, genes belonging to the “tumor necrosis alpha (TNFA) signaling via NFkB” category were the most significantly overexpressed in Braf^V600E^+K5-Tert tumors (full details in Suppl Table S7). To confirm this result, we assessed the levels of phospho-p65, a key component of the TNFA-mediated canonical NFkB signaling (37), in unselected thyroid tumors from Braf^V600E^, Braf^V600E^+Tert^-123C>T^ and Braf^V600E^+K5-Tert 20-week animals. Western blotting showed higher levels of phospho-p65 in telomerase-reactivated extracts, particularly Braf^V600E^+Tert^-123C>T^, suggesting this pathway is also overactivated in these specimens (Fig 5A and Suppl Fig 4C). Overall, RNAseq unveiled that key mediators of inflammation, likely in crosstalk with the tumor immune infiltrate, may play a role in Tert-induced thyroid cancer progression.

**Figure 5.**
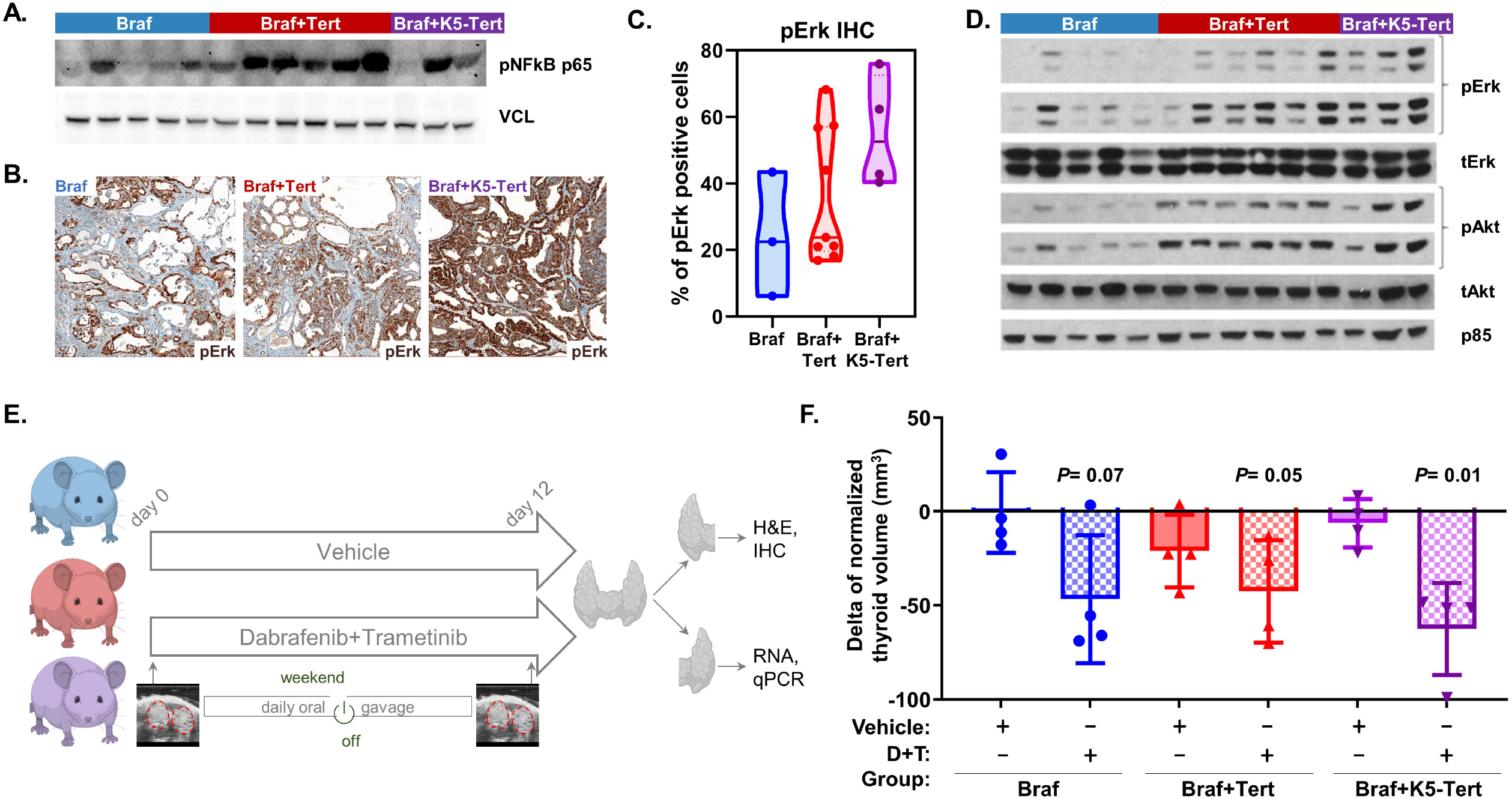
Validation of telomerase-reactivated pathways and consequences for treatment. **A**. Western blot using protein extracts from thyroid tumors from 20-week-old animals with the indicated genotypes for phospho-NFkB p65 and vinculin (VCL, loading control). **B**. Representative images from phospho-Erk (pErk) immunohistochemistry performed in mouse tumors from 20-week-old animals with the indicated genotypes. **C**. Quantification of the percentage of tumor cells staining positive for pErk in mice with the indicated genotypes. **D**. Western blot using protein extracts from thyroid tumors from 20-week-old animals with the indicated genotypes for phospho- and total Erk (MAPK pathway) and Akt (PI3K effector), as well as p85 (loading control). **E**. Schematic representation of the **in vivo** treatment of mouse models. Twenty-four 20-week animals from the Braf^V600E^, Braf^V600E^+Tert^-123C>T^ and Braf^V600E^+K5-Tert groups were randomly assigned and treated with either vehicle or dabrafenib plus tramentinib for 12 days. Thyroid volume was assessed by ultrasound on days 1 and 12. All mice were sacrificed on day 12, their thyroids were harvested, and each thyroid lobe was processed for either hematoxylin & eosin (H&E) staining and immunohistochemistry (IHC) or digested and preserved in Trizol for RNA extraction and subsequent real-time quantitative PCR (qPCR) assays. **F**. Thyroid tumor volumes, measured by ultrasound, in animals with the indicated genotypes and treatment groups. Results are expressed as the delta of normalized thyroid volumes between day 12 (last day of the experiment) and day 1 (first day). Abbreviations: “Braf+Tert” and “Braf+K5-Tert” labels denote “Braf^V600E^+Tert^-123C>T^” and “Braf^V600E^+keratin 5-driven Tert^”^ genotypes, respectively; WT= wildtype; YFP= yellow fluorescent protein; D+T= Dabrafenib plus Trametinib.

In addition, pathway analysis showed that several *bona fide* signaling hubs in thyroid cancer pathogenesis, including the MAPK and PI3K/AKT pathways, ranked high as hyperactivated pathways in Braf^V600E^+K5-Tert tumors (Fig 4F). Despite some inter-mouse variability, MAPK overactivation was confirmed via IHC analysis for phospho-Erk (pErk) of tumors from 20-week animals, showing that, compared to Braf^V600E^, Braf^V600E^+K5-Tert have higher proportion of tumor cells staining positive for pErk (***P***= 0.0673, Fig 5B-C). Western blotting with protein extracts from 20-week mouse tumors from each genotype further showed increased phosphorylation levels for Erk and Akt in Tert-engineered tumors (Fig 5D and Suppl Fig S4D). Because MAPK pathway activation in human and mouse thyroid tumors inversely correlate with transcriptional programs of thyroid differentiation, we evaluated whether key thyroid transcription factors and iodine metabolism genes were suppressed across different genotypes in our RNAseq data. In general, thyroid transcription factors *Pax8, Nkx2-1* and *Foxe1*, and iodine metabolism genes *Tshr, Slc5a5* and *Tpo*, tended to be reduced in Braf^V600E^+K5-Tert cells, which showed the highest MAPK overactivation, whereas other than for *Tshr* levels, we did not find suppressed markers of thyroid differentiation in Braf^V600E^+Tert^-123C>T^ tumors (Suppl Fig S4E).

### Thyroid tumors with telomerase reactivation remain responsive to MAPK inhibition treatment

Since MAPK pathway overactivation was detected in tumors with high telomerase levels, we evaluated whether *in vivo* treatment with dabrafenib (Raf inhibitor) plus trametinib (Mek inhibitor) impacted tumor growth (Fig 5E). Sustained treatment for 12 days was well tolerated, and body weight of treated mice did not vary over time (Suppl Fig S5A). Compared to vehicle, dabrafenib plus trametinib treatment reduced tumor volume in all three groups (Braf^V600E^, Braf^V600E^+Tert^-123C>T^ and Braf^V600E^+K5-Tert) (Fig 5F and Suppl Fig S5B). Thyroid differentiation markers increased upon MAPK inhibition in all three genotype groups, but both models with telomerase reactivation showed slightly lower re-expression of these genes, compared to Braf^V600E^ animals, perhaps reflecting their lesser baseline mRNA levels for thyroid genes (Suppl Fig S5D). Overall, despite the observed higher MAPK output of telomerase-reactivated thyroid tumors, combination treatment with dabrafenib plus trametinib was effective at reducing tumor volume across all evaluated mouse models.

## DISCUSSION

Telomerase reactivation occurs in 90% of tumors and is considered a hallmark of cancer (38,39). The discovery of *TERT* promoter mutations (TPMs) occurring in multiple cancers, and the fact that TPMs are more prevalent in metastatic/advanced forms of the disease, have heightened interest in studying this oncogene. Accurate experimental models to mechanistically assess the biological consequences of telomerase reactivation in a neoplastic background are scarce. Here we studied how telomerase re-expression contributes to thyroid cancer progression in Braf^V600E^-driven murine tumors. To this end, we edited a C>T change in the mouse Tert promoter (Tert^-123C>T^), which we believe is the orthologue nucleotide of human TERT^-124C>T^, and also employed a previously generated model of Tert overexpression (K5-Tert). Our experiments showed that Tert upregulation cooperates with oncogenic Braf to promote advanced thyroid cancers, which was associated with activation of oncogenic pathways unrelated to telomere maintenance. Tert^-123C>T^ mice were born at Mendelian rates comparable to Tert wildtype littermates and displayed no cancer phenotypes, consistent with the observation that this promoter variant is found in families that have no abnormalities other than a higher frequency of metastatic melanoma (1).

We and others have previously reported that TPMs are enriched in advanced thyroid tumors, i.e., PDTC and ATC (4,5). Furthermore, in the context of constitutive MAPK pathway activation, they are predictors of poorer outcomes within each subtype. The presence of TPMs in combination with BRAF^V600E^ is associated with more aggressive PTCs, including higher mortality rates (9,40,41). In advanced disease, TPMs correlate with the presence of distant metastases in PDTC and with diminished survival in ATC patients (13). This is particularly relevant in thyroid cancers, which are genetically simple tumors that remain exquisitely dependent on MAPK activation, making them good models to study MAPK-TERT contributions to cancer biology. The models described here could thus be useful tools to determine the cellular processes by which Braf-driven clones, upon the reactivation of Tert, evolve to more aggressive and less differentiated cancers.

Our *in vivo* experiments showed that Tert upregulation, regardless of its mechanism of overexpression, induced Braf^V600E^-driven PTCs to progress to PDTC/HGDTC. This phenomenon was observed starting at 20 weeks, when no Braf^V600E^ animals, but around one-third of Braf^V600E^+Tert^-123C>T^ and Braf^V600E^+K5-Tert mice showed advanced phenotypes. The variability in Tert-driven thyroid cancer progression observed in our models remains to be fully understood: it is unclear, at this point, whether thyroid dedifferentiation is a stochastic process or follows a defined pattern of events. In addition, a subset of older Braf^V600E^+Tert^-123C>T^, but none of the Braf^V600E^ group, displayed ATC-like features. These included a strong infiltration of the tumor microenvironment by myeloid cells, particularly M2-macrophages and myeloid-derived suppressor cells (MDSC). This is relevant because human ATCs are known to be infiltrated by tumor-promoting macrophages and MDSCs (42-44). Given that BRAF^V600E^+TERT^-124C>T^ is the most common genetic combination in ATC (13), our findings suggest that telomerase reactivation is a stepping stone towards the progressive immunosuppression that tracks with PTC-to-ATC transformation (45). In any case, some of the pathways that we identified to be activated in murine thyroid tumor cells upon Tert upregulation (see next paragraphs) suggest a continuous crosstalk between cancer cells and components of the tumor immune milieu.

Another feature of these mouse models that conveys potentially important translational consequences is the increased activation of Braf^V600E^-induced MAPK signaling which we observed in telomerase-upregulated tumors. The MAPK pathway is a central signaling hub in both PTC and PDTC/ATC. The use of dabrafenib (RAF inhibitor) plus trametinib (MEK inhibitor) in ATCs harboring BRAF^V600E^ mutations remains one of the few effective therapeutic strategies in these otherwise highly lethal cancers (46). Our *in vivo* treatment with dabrafenib plus trametinib showed comparable tumor volume reductions in telomerase-upregulated specimens. The relatively small number of animals per group and condition was insufficient for robust statistical association; however, genotype-dependent insights in response to MAPK treatment were hinted. Overall, future work using these and other models, combined with clinical trials with extensive genomic characterization, might ultimately help refine the specifics of MAPK blockade in ATC patients, given that most of their tumors harbor TPMs. Of note, interactions between TERT and MAPK pathway effectors have been previously reported in other tumors (47).

We observed heterogeneity in Tert re-expression in Braf^V600E^+Tert^-123C>T^ tumors, in line with reports showing intermittent expression of TERT in human cancer cells (48). A much more granular evaluation of promoter binding preferences around the mouse *Tert* promoter would be needed to unveil these processes, which are likely related with chromatin accessibility, histone modifications and/or differential binding of transcriptions factors at the *Tert* locus. In this regard, Tert^-123C>T^ created a consensus binding sequence for transcription factors of the ETS (“E26 transformation specific”) family which was identical to the one generated by TERT^-124C>T^ in humans (Fig. 1A). The ETS family is a diverse group of proteins, and the specific factor(s) influencing transcription from the *TERT* mutant promoter in thyroid cancers remain to be fully elucidated (49-52). In addition, MAPK activation is also known to control *TERT* transcription (52-54). Overall, although these studies are beyond the scope of this paper, we believe that the Braf^V600E^+Tert^-123C>T^ model can be leveraged for future work on *Tert* mutant promoter regulation. The ease at generating tumors and deriving cell lines from these animals should pave the way towards this goal. Of note, despite the intra-tumoral heterogeneity in Tert expression, cell lines derived from these tumors maintained a genotype-dependent increase in Tert transcription, which might reflect a selection process by which cells expressing higher levels of Tert are favored over time.

We acknowledge that mouse models are not ideal settings to study the canonical role of telomerase biology in cancer, i.e., telomere maintenance. This limitation has been extensively reported in the literature and relies on two features of mouse cells which do not occur in humans: having longer telomeres and retaining some degree of telomerase expression in their adult tissues (21,22). Despite these, here we demonstrate that Tert reactivation leads to thyroid cancer progression and upregulation of non-telomeric pathways. Indeed, the demonstration that telomerase upregulation causes disease progression in mice further substantiates the evidence that non-canonical effects of Tert play an important role in this process. Our RNAseq-derived pathway analyses support a role for cytokines and chemokines produced by Tert-activated thyroid cancer cells, probably in response to signals from the tumor microenvironment. Signaling of these molecules via the canonical (i.e., TNFA-activated) NFkB pathway, a key node linking inflammation, immune-related effects and cancer, was particularly enriched in our analysis. Inhibition of the NFkB/TNFA signaling in human thyroid cancer cell lines, most of which are now known to harbor *TERT* promoter mutations (55), has been shown to impact cell proliferation and invasion (56). Interestingly, our results in murine models recapitulate observations reported in human cancers, i.e., telomerase controlling NFkB-dependent transcriptional programs, and NFkB in turn determining TERT nuclear localization and influencing transcription from the mutant *TERT* promoter, in what was described as a feed-forward mechanism (57-59).

Tert^-123C>T^ constitutes the first attempt at engineering a promoter mutation in a murine *Tert* non-coding locus. To our knowledge, there had not been *in vivo* studies of TPMs in genetically engineered mouse models. The closest effort is a recent publication in which human stem cells were engineered for TERT^-124C>T^ mutation, differentiated into neural precursors and orthotopically injected into mice to study glioblastoma in the context of *EGFR* overexpression and loss of *CDKN2A* and *PTEN* (60). Our results showing that murine Tert reactivation promotes cancer progression in oncogenic Braf-driven thyroid tumors, open the door at devising equivalent studies in tumors from other lineages with high prevalence of TPMs. Although some of the effects reported here might pertain exclusively to thyroid tumors, we believe that the combination of the Tert^-123C>T^ allele with, for instance, tissue-specific drivers of melanoma or glioblastoma, such as the Tyr:NRas^Q61K^/INK4a^-/-^ (61,62) or EGFRvIII (63) mouse models, respectively, can provide hints into the telomerase-dependent biology of those cancers. Of note, although the K5-Tert model was primarily used as a positive control of continuous Tert overexpression, in light of our results, we argue that employing Braf^V600E^+K5-Tert animals can be an informative setting to study telomerase-mediated thyroid cancer progression. This statement relies on the fact that the observed increase of Tert expression in Braf^V600E^+K5-Tert thyroid tumor cells was about 10-fold, in contrast to a much higher (and thus less biologically accurate) increases in other epithelial cells. It is also worth noting that other mechanisms of ***TERT*** upregulation, such as *TERT* copy number gains and aberrant methylation of the *TERT* promoter, have been identified in advanced thyroid cancers (33,64,65).

In conclusion, here we show that the *in vivo* engineering of a non-coding alteration in the mouse *Tert* promoter induces thyroid cancer progression. We show that Tert upregulation induces changes in the transcriptome of fluorescence-sorted cancer cells isolated from murine thyroid tumors, and point to several cellular processes, not involved in telomere elongation, that are altered in Braf+Tert-driven advanced thyroid cancers. Our study can shed light into the multiple mechanistic underpinnings that telomerase reactivation bears in cancer.

## ACKNOWLEDGEMENTS

We would like to thank Talia Gebhard from the Landa lab, and Joe Giacalone from the MSKCC Mouse Genetics Core Facility, for technical assistance. We thank the Harvard Medical School Neurobiology Imaging Facility for technical assistance, as well as the Dana-Farber/Harvard Cancer Center in Boston, MA, for the use of the Specialized Histopathology Core, which provided histology and immunohistochemistry service. The MSKCC Mouse Genetics Core Facility is supported by the NCI Cancer Center Support Grant P30 CA008748. The Dana-Farber/Harvard Cancer Center is supported in part by an NCI Cancer Center Support Grant # NIH 5 P30 CA06516.

## REFERENCES

1. Horn S, Figl A, Rachakonda PS, Fischer C, Sucker A, Gast A, et al. TERT promoter mutations in familial and sporadic melanoma. Science 2013;339:959–61

2. Huang FW, Hodis E, Xu MJ, Kryukov GV, Chin L, Garraway LA. Highly recurrent TERT promoter mutations in human melanoma. Science 2013;339:957–9

3. Killela PJ, Reitman ZJ, Jiao Y, Bettegowda C, Agrawal N, Diaz LA, Jr., et al. TERT promoter mutations occur frequently in gliomas and a subset of tumors derived from cells with low rates of self-renewal. Proc Natl Acad Sci U S A 2013;110:6021–6

4. Landa I, Ganly I, Chan TA, Mitsutake N, Matsuse M, Ibrahimpasic T, et al. Frequent somatic TERT promoter mutations in thyroid cancer: higher prevalence in advanced forms of the disease. J Clin Endocrinol Metab 2013;98:E1562–6

5. Liu X, Bishop J, Shan Y, Pai S, Liu D, Murugan AK, et al. Highly prevalent TERT promoter mutations in aggressive thyroid cancers. Endocr Relat Cancer 2013;20:603–10

6. Weinhold N, Jacobsen A, Schultz N, Sander C, Lee W. Genome-wide analysis of noncoding regulatory mutations in cancer. Nat Genet 2014;46:1160–5

7. Fredriksson NJ, Ny L, Nilsson JA, Larsson E. Systematic analysis of noncoding somatic mutations and gene expression alterations across 14 tumor types. Nat Genet 2014;46:1258–63

8. Rheinbay E, Nielsen MM, Abascal F, Wala JA, Shapira O, Tiao G, et al. Analyses of non-coding somatic drivers in 2,658 cancer whole genomes. Nature 2020;578:102–11

9. Melo M, da Rocha AG, Vinagre J, Batista R, Peixoto J, Tavares C, et al. TERT promoter mutations are a major indicator of poor outcome in differentiated thyroid carcinomas. J Clin Endocrinol Metab 2014;99:E754–65

10. Griewank KG, Murali R, Puig-Butille JA, Schilling B, Livingstone E, Potrony M, et al. TERT promoter mutation status as an independent prognostic factor in cutaneous melanoma. J Natl Cancer Inst 2014;106

11. Blasco MA. Telomeres and human disease: ageing, cancer and beyond. Nat Rev Genet 2005;6:611–22

12. Martinez P, Blasco MA. Telomeric and extra-telomeric roles for telomerase and the telomere-binding proteins. Nat Rev Cancer 2011;11:161–76

13. Landa I, Ibrahimpasic T, Boucai L, Sinha R, Knauf JA, Shah RH, et al. Genomic and transcriptomic hallmarks of poorly differentiated and anaplastic thyroid cancers. J Clin Invest 2016;126:1052–66

14. Pozdeyev N, Gay L, Sokol ES, Hartmaier RJ, Deaver KE, Davis SN, et al. Genetic analysis of 779 advanced differentiated and anaplastic thyroid cancers. Clin Cancer Res 2018

15. Shi X, Liu R, Qu S, Zhu G, Bishop J, Liu X, et al. Association of TERT promoter mutation 1,295,228 C>T with BRAF V600E mutation, older patient age, and distant metastasis in anaplastic thyroid cancer. J Clin Endocrinol Metab 2015;100:E632–7

16. Cancer Genome Atlas Research N. Integrated genomic characterization of papillary thyroid carcinoma. Cell 2014;159:676–90

17. Gonzalez-Suarez E, Samper E, Ramirez A, Flores JM, Martin-Caballero J, Jorcano JL, et al. Increased epidermal tumors and increased skin wound healing in transgenic mice overexpressing the catalytic subunit of telomerase, mTERT, in basal keratinocytes. EMBO J 2001;20:2619–30

18. Artandi SE, Alson S, Tietze MK, Sharpless NE, Ye S, Greenberg RA, et al. Constitutive telomerase expression promotes mammary carcinomas in aging mice. Proc Natl Acad Sci U S A 2002;99:8191–6

19. Canela A, Martin-Caballero J, Flores JM, Blasco MA. Constitutive expression of tert in thymocytes leads to increased incidence and dissemination of T-cell lymphoma in Lck-Tert mice. Mol Cell Biol 2004;24:4275–93

20. Gonzalez-Suarez E, Geserick C, Flores JM, Blasco MA. Antagonistic effects of telomerase on cancer and aging in K5-mTert transgenic mice. Oncogene 2005;24:2256–70

21. Prowse KR, Greider CW. Developmental and tissue-specific regulation of mouse telomerase and telomere length. Proc Natl Acad Sci U S A 1995;92:4818–22

22. Kipling D, Cooke HJ. Hypervariable ultra-long telomeres in mice. Nature 1990;347:400–2

23. Pinello L, Canver MC, Hoban MD, Orkin SH, Kohn DB, Bauer DE, et al. Analyzing CRISPR genomeediting experiments with CRISPResso. Nature biotechnology 2016;34:695–7

24. Franco AT, Malaguarnera R, Refetoff S, Liao XH, Lundsmith E, Kimura S, et al. Thyrotrophin receptor signaling dependence of Braf-induced thyroid tumor initiation in mice. Proc Natl Acad Sci U S A 2011;108:1615–20

25. Saqcena M, Leandro-Garcia LJ, Maag JLV, Tchekmedyian V, Krishnamoorthy GP, Tamarapu PP, et al. SWI/SNF Complex Mutations Promote Thyroid Tumor Progression and Insensitivity to Redifferentiation Therapies. Cancer Discov 2021;11:1158–75

26. Bankhead P, Loughrey MB, Fernandez JA, Dombrowski Y, McArt DG, Dunne PD, et al. QuPath: Open source software for digital pathology image analysis. Sci Rep 2017;7:16878

27. Cornwell M, Vangala M, Taing L, Herbert Z, Koster J, Li B, et al. VIPER: Visualization Pipeline for RNA-seq, a Snakemake workflow for efficient and complete RNA-seq analysis. BMC Bioinformatics 2018;19:135

28. Schneider CA, Rasband WS, Eliceiri KW. NIH Image to ImageJ: 25 years of image analysis. Nat Methods 2012;9:671–5

29. Callicott RJ, Womack JE. Real-time PCR assay for measurement of mouse telomeres. Comp Med 2006;56:17–22

30. Wei GH, Badis G, Berger MF, Kivioja T, Palin K, Enge M, et al. Genome-wide analysis of ETS-family DNA-binding in vitro and in vivo. EMBO J 2010;29:2147–60

31. Bullock M, Ren Y, O’Neill C, Gill A, Aniss A, Sywak M, et al. TERT promoter mutations are a major indicator of recurrence and death due to papillary thyroid carcinomas. Clin Endocrinol (Oxf) 2016;85:283–90

32. Vinagre J, Almeida A, Populo H, Batista R, Lyra J, Pinto V, et al. Frequency of TERT promoter mutations in human cancers. Nature communications 2013;4:2185

33. Montero-Conde C, Leandro-Garcia LJ, Martinez-Montes AM, Martinez P, Moya FJ, Leton R, et al. Comprehensive molecular analysis of immortalization hallmarks in thyroid cancer reveals new prognostic markers. Clin Transl Med 2022;12:e1001

34. Hiltzik D, Carlson DL, Tuttle RM, Chuai S, Ishill N, Shaha A, et al. Poorly differentiated thyroid carcinomas defined on the basis of mitosis and necrosis: a clinicopathologic study of 58 patients. Cancer 2006;106:1286–95

35. Volante M, Collini P, Nikiforov YE, Sakamoto A, Kakudo K, Katoh R, et al. Poorly differentiated thyroid carcinoma: the Turin proposal for the use of uniform diagnostic criteria and an algorithmic diagnostic approach. Am J Surg Pathol 2007;31:1256–64

36. Baloch ZW, Asa SL, Barletta JA, Ghossein RA, Juhlin CC, Jung CK, et al. Overview of the 2022 WHO Classification of Thyroid Neoplasms. Endocr Pathol 2022;33:27–63

37. Taniguchi K, Karin M. NF-kappaB, inflammation, immunity and cancer: coming of age. Nat Rev Immunol 2018;18:309–24

38. Kim NW, Piatyszek MA, Prowse KR, Harley CB, West MD, Ho PL, et al. Specific association of human telomerase activity with immortal cells and cancer. Science 1994;266:2011–5

39. Hanahan D, Weinberg RA. Hallmarks of cancer: the next generation. Cell 2011;144:646–74

40. Liu R, Bishop J, Zhu G, Zhang T, Ladenson PW, Xing M. Mortality Risk Stratification by Combining BRAF V600E and TERT Promoter Mutations in Papillary Thyroid Cancer: Genetic Duet of BRAF and TERT Promoter Mutations in Thyroid Cancer Mortality. JAMA Oncol 2017;3:202–8

41. Xing M, Liu R, Liu X, Murugan AK, Zhu G, Zeiger MA, et al. BRAF V600E and TERT promoter mutations cooperatively identify the most aggressive papillary thyroid cancer with highest recurrence. J Clin Oncol 2014;32:2718–26

42. Caillou B, Talbot M, Weyemi U, Pioche-Durieu C, Al Ghuzlan A, Bidart JM, et al. Tumorassociated macrophages (TAMs) form an interconnected cellular supportive network in anaplastic thyroid carcinoma. PLoS One 2011;6:e22567

43. Ryder M, Ghossein RA, Ricarte-Filho JC, Knauf JA, Fagin JA. Increased density of tumorassociated macrophages is associated with decreased survival in advanced thyroid cancer. Endocr Relat Cancer 2008;15:1069–74

44. Suzuki S, Shibata M, Gonda K, Kanke Y, Ashizawa M, Ujiie D, et al. Immunosuppression involving increased myeloid-derived suppressor cell levels, systemic inflammation and hypoalbuminemia are present in patients with anaplastic thyroid cancer. Mol Clin Oncol 2013;1:959–64

45. Luo H, Xia X, Kim GD, Liu Y, Xue Z, Zhang L, et al. Characterizing dedifferentiation of thyroid cancer by integrated analysis. Sci Adv 2021;7

46. Subbiah V, Kreitman RJ, Wainberg ZA, Cho JY, Schellens JHM, Soria JC, et al. Dabrafenib and Trametinib Treatment in Patients With Locally Advanced or Metastatic BRAF V600-Mutant Anaplastic Thyroid Cancer. J Clin Oncol 2018;36:7–13

47. Li Z, Ivanov AA, Su R, Gonzalez-Pecchi V, Qi Q, Liu S, et al. The OncoPPi network of cancer-focused protein-protein interactions to inform biological insights and therapeutic strategies. Nature communications 2017;8:14356

48. Ravindranathan A, Cimini B, Diolaiti ME, Stohr BA. Preliminary development of an assay for detection of TERT expression, telomere length, and telomere elongation in single cells. PLoS One 2018;13:e0206525

49. Liu R, Zhang T, Zhu G, Xing M. Regulation of mutant TERT by BRAF V600E/MAP kinase pathway through FOS/GABP in human cancer. Nature communications 2018;9:579

50. Bullock M, Lim G, Zhu Y, Aberg H, Kurdyukov S, Clifton-Bligh R. ETS Factor ETV5 Activates the Mutant Telomerase Reverse Transcriptase Promoter in Thyroid Cancer. Thyroid 2019;29:1623–33

51. Song YS, Yoo SK, Kim HH, Jung G, Oh AR, Cha JY, et al. Interaction of BRAF-induced ETS factors with mutant TERT promoter in papillary thyroid cancer. Endocr Relat Cancer 2019;26:629–41

52. Thornton CEM, Hao J, Tamarapu PP, Landa I. Multiple ETS Factors Participate in the Transcriptional Control of TERT Mutant Promoter in Thyroid Cancers. Cancers (Basel) 2022;14

53. Greenberg RA, O’Hagan RC, Deng H, Xiao Q, Hann SR, Adams RR, et al. Telomerase reverse transcriptase gene is a direct target of c-Myc but is not functionally equivalent in cellular transformation. Oncogene 1999;18:1219–26

54. Takakura M, Kyo S, Kanaya T, Hirano H, Takeda J, Yutsudo M, et al. Cloning of human telomerase catalytic subunit (hTERT) gene promoter and identification of proximal core promoter sequences essential for transcriptional activation in immortalized and cancer cells. Cancer Res 1999;59:551–7

55. Landa I, Pozdeyev N, Korch C, Marlow LA, Smallridge RC, Copland JA, et al. Comprehensive genetic characterization of human thyroid cancer cell lines: a validated panel for preclinical studies. Clin Cancer Res 2019

56. Bauerle KT, Schweppe RE, Haugen BR. Inhibition of nuclear factor-kappa B differentially affects thyroid cancer cell growth, apoptosis, and invasion. Mol Cancer 2010;9:117

57. Li Y, Zhou QL, Sun W, Chandrasekharan P, Cheng HS, Ying Z, et al. Non-canonical NF-kappaB signalling and ETS1/2 cooperatively drive C250T mutant TERT promoter activation. Nature cell biology 2015;17:1327–38

58. Ghosh A, Saginc G, Leow SC, Khattar E, Shin EM, Yan TD, et al. Telomerase directly regulates NF-kappaB-dependent transcription. Nature cell biology 2012;14:1270–81

59. Akiyama M, Hideshima T, Hayashi T, Tai YT, Mitsiades CS, Mitsiades N, et al. Nuclear factor-kappaB p65 mediates tumor necrosis factor alpha-induced nuclear translocation of telomerase reverse transcriptase protein. Cancer Res 2003;63:18–21

60. Miki S, Koga T, McKinney AM, Parisian AD, Tadokoro T, Vadla R, et al. TERT promoter C228T mutation in neural progenitors confers growth advantage following telomere shortening in vivo. Neuro Oncol 2022;24:2063–75

61. Ackermann J, Frutschi M, Kaloulis K, McKee T, Trumpp A, Beermann F. Metastasizing melanoma formation caused by expression of activated N-RasQ61K on an INK4a-deficient background. Cancer Res 2005;65:4005–11

62. Sharpless NE, Kannan K, Xu J, Bosenberg MW, Chin L. Both products of the mouse Ink4a/Arf locus suppress melanoma formation in vivo. Oncogene 2003;22:5055–9

63. Stockhausen MT, Broholm H, Villingshoj M, Kirchhoff M, Gerdes T, Kristoffersen K, et al. Maintenance of EGFR and EGFRvIII expressions in an in vivo and in vitro model of human glioblastoma multiforme. Exp Cell Res 2011;317:1513–26

64. McKelvey BA, Zeiger MA, Umbricht CB. Characterization of TERT and BRAF copy number variation in papillary thyroid carcinoma: An analysis of the cancer genome atlas study. Genes Chromosomes Cancer 2021;60:403–9

65. Lee DD, Leao R, Komosa M, Gallo M, Zhang CH, Lipman T, et al. DNA hypermethylation within TERT promoter upregulates TERT expression in cancer. J Clin Invest 2019;129:223–9

66. Mercer K, Giblett S, Green S, Lloyd D, DaRocha Dias S, Plumb M, et al. Expression of endogenous oncogenic V600EB-raf induces proliferation and developmental defects in mice and transformation of primary fibroblasts. Cancer Res 2005;65:11493–500

