## Supplementary Figure and Table Legends for "Telomerase reactivation induces progression of mouse Braf^V600E^-driven thyroid cancers without telomere lengthening"

**SUPPLEMENTARY FIGURE LEGENDS**

**Supplementary Figure S1. Pairwise alignment of human:mouse telomerase reverse transcriptase (*TERT*) gene sequences.** Full alignment of human *vs.* mouse *TERT* sequences around the transcription start site (ATG, highlighted in yellow), including a ~300bp portion of the proximal promoter. The following nucleotides are highlighted for human *TERT*: c.-146C (light blue) and c.-124C (red). Mouse *Tert* c.-123C is highlighted in green and aligns well with human *TERT* c.-124C. Results from the Needle algorithm, available at the European Bioinformatics Institute website (EMBL-EBI; https://www.ebi.ac.uk/Tools/psa/), are shown.

**Supplementary Figure S2. Additional features of Braf and Braf+Tert murine tumors. A.** Thyroid tumors generated in 10-week-old Braf^V600E^-mutant animals displaying the defining characteristics of papillary thyroid cancers (PTC): presence of papillae (green arrowheads) and nuclear features such as nuclear membrane irregularity (in grey), chromatin margination (blue), nuclear grooves (yellow) and nuclear pseudoinclusion (orange). **B.** Examples of 20-week Braf^V600E^+Tert^-123C>T^ thyroid tumors with PDTC phenotypes, represented by areas of solid growth which correlate with highly proliferative cells (as per Ki67 immunostaining) and with abundance of mitotic figures (yellow arrowheads). **C.** Thyroid volume, assessed by ultrasound, of mouse thyroid tumors of 16-to-20-week animals with the indicated genotypes. Tumors in which Tert was reactivated showed a non-significant trend towards larger volumes: t-test associated *P* values 0.18 and 0.14 for the Braf^V600E^ *vs.* Braf^V600E^+Tert^-123C>T^ and the Braf *vs.* Braf^V600E^+K5-Tert comparisons, respectively. **D.** Kaplan-Meier curves showing survival of tumor-bearing Braf^V600E^ *vs.* Braf^V600E^+Tert^-123C>T^ animals at 30 weeks. **E.** Examples of 40-week Braf^V600E^ (top panel) and Braf^V600E^+Tert^-123C>T^ tumors (middle and bottom panels). Some Braf^V600E^ animals at this age develop signs of PDTC transformation (top panel) and typically retain Pax8 expression (top, right). Only Braf^V600E^+Tert^-123C>T^ animals of this age developed tumors with anaplastic thyroid cancer (ATC)-like features within a PTC component (middle and bottom panels), overall loss of Pax8 with spindle cells retaining Pax8 positivity (middle, right), spindle cells (bottom, center, yellow arrows) and positivity for F4/80 stain, denoting macrophage infiltration (bottom, right); **F.** Example of a 20-week Braf^V600E^+K5-Tert tumor with similar features than tumors from panel E, i.e., ATC-like areas, spindle cells (yellow arrowhead) with focal positivity for Pax8 (spindle cells with red arrowheads). **G.** Representative examples of immunostaining performed on thyroid specimens from 30-to-40 week-old-animals with the indicated genotypes and the following antibodies: Cd11b (pan-myeloid stain, left), F4/80 (macrophages, middle) and Arg1 (M2-like). **H.** Quantification of Cd11b, F4/80 and Arg1 stains in 30-to-40 week-old-animals. Each dot represents an independent animal, and results are shown as median and interquartile range. Abbreviations: “Braf+Tert” and “Braf+K5-Tert” labels denote “Braf^V600E^+Tert^-123C>T^” and “Braf^V600E^+keratin 5-driven Tert^”^ genotypes, respectively. WT=wildtype.

**Supplementary Figure S3. Analysis of intratumoral heterogeneity of Tert transcription via RNAscope.** Representative examples of tumors from 20-week animals. **A-F.** Braf^V600E^+Tert^-123C>T^ mouse showing a poorly differentiated thyroid cancer (PDTC); **G-L.** Braf^V600E^ mouse with a papillary thyroid cancer (PTC). Hematoxylin-eosin (H&E) staining and RNAscope detection of Tert transcripts (green) are displayed side-by-side, and dotted boxes show zoomed-in views of specific regions within each tumor. DAPI staining (in blue) is shown for contrast. The PDTC area of the Braf^V600E^+Tert^-123C>T^ tumor (**C**) correlates with higher fluorescent signals, indicating increased Tert transcription within cells (**F**). In contrast, a PTC area within the same tumor (**A**) shows fewer fluorescent dots (**D**). Braf^V600E^ mice are homogeneous tumors displaying PTC phenotypes in all regions (**G, I**), which correlate with similar fluorescent signals (**J, L**) which are typically lower than those in Braf^V600E^+Tert^-123C>T^ tumors.

**Supplementary Figure S4. Additional analyses of RNA sequencing data from mouse tumors with telomerase reactivation. A.** Unsupervised hierarchical clustering of RNAseq data in mouse tumors with the indicated genotypes. **B.** KEGG (Kyoto Encyclopedia of Genes and Genomes) pathway analysis of RNAseq data showing the top upregulated terms in Braf^V600E^+Tert^-123C>T^ tumors, compared to Braf^V600E^, from 20-week mice. Results are based on unadjusted p-values from this comparison, and non-significant (p-value>0.05) are shown as grey bars. **C.** Quantification of phospho-NFkB p65 band intensity from western blot from figure 5A. Values are normalized with intensities from their respective loading control. Braf^V600E^ tumors are used as baseline, and p-values for Braf^V600E^+Tert^-123C>T^ and Braf^V600E^+K5-Tert are 0.0087 and 0.1429, respectively. **D.** Quantification of phospho-Erk (left panel) and phospho-Akt (right) band intensities from western blot from figure 5D. Values are normalized with intensities from their respective loading controls. Braf^V600E^ tumors are used as baseline, and p-values for pErk levels for Braf^V600E^+Tert^-123C>T^ and Braf^V600E^+K5-Tert are 0.3101 and 0.0673, respectively. P-values for pAkt for Braf^V600E^+Tert^-123C>T^ and Braf^V600E^+K5-Tert are 0.0002 and 0.0245, respectively. **E.** RNAseq normalized counts for genes involved in thyroid differentiation and iodine metabolism from mouse tumors with the indicated genotypes. Asterisks represent significant p-values for each condition, using Braf^V600E^ as control. Abbreviations: “Braf+Tert” and “Braf+K5-Tert” labels denote “Braf^V600E^+Tert^-123C>T^” and “Braf^V600E^+keratin 5-driven Tert^”^ genotypes, respectively.

**Supplementary Figure S5. Detailed information of the *in vivo* dabrafenib plus trametinib treatment experiment. A.** Evolution of mouse body weight, expressed in grams (g) of all animals included in the study: 12 treated with vehicle (left panel) and 12 treated with the dabrafenib plus trametinib combo (right). Each mouse is color-coded by genotype, as indicated. Some lines are not visible due to overlaps of identical weights. **B.** Hematoxylin & eosin (H&E) staining of all 24 animals included in this study, distributed by genotype and treatment group, as indicated. **C.** Phospho-Erk (pErk) staining of all 24 animals included in this study, distributed by genotype and treatment group, as indicated. **D.** Expression levels, assessed by qPCR, for thyroid-specific genes *Slc5a5* (also known as *Nis*), *Pax8* and *Tpo*, in mouse tumors from the indicated genotypes and treatment groups. Results are expressed in relative units (r.u.) and are normalized by b-actin (housekeeping gene) for each sample. Abbreviations: “Braf+Tert” and “Braf+K5-Tert” labels denote “Braf^V600E^+Tert^-123C>T^” and “Braf^V600E^+keratin 5-driven Tert^”^ genotypes, respectively.

**SUPPLEMENTARY TABLE LEGENDS**

**Supplementary Table S1.** List of primers used for quantitative PCR.

**Supplementary Table S2.** CRISPRESSO sequencing results for an F0 animal in which the Tert c.-123C>T (red font) mutation was engineered via CRISPR/Cas9 editing. The second row represents the desired on-target result without off-target effects.

**Supplementary Table S3.** Differentially expressed genes from RNAseq data when comparing Braf^V600E^ *vs.* Braf^V600E^+Tert^-123C>T^ tumors from 20-week animals. Positive fold change values (log2FoldChange) represent genes overexpressed in the Braf^V600E^+Tert^-123C>T^ group.

**Supplementary Table S4.** Differentially expressed genes from RNAseq data when comparing Braf^V600E^ *vs.* Braf^V600E^+K5-Tert tumors from 20-week animals. Positive fold change values (log2FoldChange) represent genes overexpressed in the Braf^V600E^+K5-Tert group.

**Supplementary Table S5.** Full list of KEGG (Kyoto Encyclopedia of Genes and Genomes) pathway analysis “up terms” for the Braf^V600E^ *vs.* Braf^V600E^+K5-Tert comparison.

**Supplementary Table S6.** Full list of KEGG (Kyoto Encyclopedia of Genes and Genomes) pathway analysis “up terms” for the Braf^V600E^ *vs.* Braf^V600E^+Tert^-123C>T^ comparison.

**Supplementary Table S7.** Full list of significantly upregulated terms (q-value<0.05) in Braf^V600E^+K5-Tert tumors, compared to Braf^V600E^, from 20-week animals, employing the GSEA (gene set enrichment analysis) hallmarks of cancer database on RNAseq data.
