## Supplementary Figures S1-S5 for "Telomerase reactivation induces progression of mouse Braf^V600E^-driven thyroid cancers without telomere lengthening"

### Suppl Fig S1

```
########################################
# Program: needle
# Commandline: needle (GLOBAL)

# Aligned_sequences: 2
# 1: Human
# 2: Mouse
# Matrix: EDNAFULL
# Gap_penalty: 10.0
# Extend_penalty: 0.5
#
# Length: 665
# Identity:     397/665 (59.7%)
# Similarity:   397/665 (59.7%)
# Gaps:         155/665 (23.3%)
# Score: 1111.0
#
```

ATG - start codon
c - TERT c.-124C
c - Tert c.-123C
c - TERT c.-146C

```
Human              1 gcgcgggcggggaagcgcggcccagacccccgggtccgcccggagcagct     50
                                |||||    ||.|||                 ||||
Mouse              1 -----------gaagc----ccgaga-----------------agca---     15

Human             51 gcgctgtcggggccaggcc---------gggctcccagtggattcgcggg     91
                      ..||||.|.||..|..||         |.||.|||..|.|.|.|.|...
Mouse             16 -ttctgtagagggaaatcctgcatgagtgcgccccctttcgttactccaa     64

Human             92 cacagacgcccaggaccgcgcttcccacgtggcggagggactggggaccc    141
                     ||||    .||||.|        .||||        .|.|||.||   ||
Mouse             65 caca----tccagca--------accac--------tgaacttgg---cc     91

Human            142 ggg-----cacc-cgtcctgccccttcacc----tt----ccagctccg-    176
                     |||     |||| .|||||   |.|.||||    ||    |||.|..||
Mouse             92 ggggaacacacctggtcct---catgcaccagcattgtgaccatcaacgg    138

Human            177 ---cctcct----ccgcgcggaccccgccccgtcc----cgacccctccc    215
                        ..|.||    |.||   |||||||||||.|||    |.||.|.|
Mouse            139 aaaagtactattgctgc---gaccccgccccttccgctacaacgctt---    182

Human            216 gggtcc-ccgg---cccagccc---cctccgggccctcccagcccctccc    258
                      ||||| ||.|   ||| ||||   ||||||    .||||||  ||||..
Mouse            183 -ggtccgcctgaatccc-gccccttcctccg----ttcccag--cctcat    224

Human            259 cttcctttc--cgcggcccc-----gccct--ctcctcgcggcgcgagtt    299
                     |||  ||||  ||.||.|.|     |.|||  .||||.||.|     .||
Mouse            225 ctt--tttcgtcgtggactctcagtggcctgggtcctggctg-----ttt    267

Human            300 tC-AGGCAGCGC---TGCGTCCTGCT--GCGCACGTGGGAAGCCC-TGGC    342
                     || |.||| |.|   |||.||.||.|  .|||||||||||.|||| |
Mouse            268 tctaagca-cacccttgcatcttggttcccgcacGTGGGAGGCCCAT---    313

Human            343 CCCGGCCACCCCCGCGATGCCGCGCGCTCCCCGCTGCCGAGCCGTGCGCT    392
                     |||||||.....|.|.|||.|.||||||||.||.||||..||.|||||||
Mouse            314 CCCGGCCTTGAGCACAATGACCCGCGCTCCTCGTTGCCCCGCGGTGCGCT    363

Human            393 CCCTGCTGCGCAGCCACTACCGCGAGGTGCTGCCGCTGGCCACGTTCGTG    442
                     |.|||||||||||||..|||||.||||||..|||||||||.||.||.|||
Mouse            364 CTCTGCTGCGCAGCCGATACCGGGAGGTGTGGCCGCTGGCAACCTTTGTG    413

Human            443 CGGCGCCTGGGGCCCCAGGGCTGGCGGCTGGTGCAGCGCGGGGACCCGGC    492
                     |||||||||||||||.|||||.|||||||.|||||.|.||||||||||..
Mouse            414 CGGCGCCTGGGGCCCGAGGGCAGGCGGCTTGTGCAACCCGGGGACCCGAA    463

Human            493 GGCTTTCCGCGCGCTGGTGGCCCAGTGCCTGGTGTGCGTGCCCTGGGACG    542
                     |...|.||||.|..||||.|||||.|||||.||||||.|||.|||||.|.
Mouse            464 GATCTACCGCACTTTGGTTGCCCAATGCCTAGTGTGCATGCACTGGGGCT    513

Human            543 CACGGCCGCCCCCCGCCGCCCCCTCCTTCCGCCAGgtgggcctcc--ccg    590
                     |||.|||.||.||.||||.||..|||||||.||||||||||||||  .||
Mouse            514 CACAGCCTCCACCTGCCGACCTTTCCTTCCACCAGgtgggcctccaggcg    563

Human            591 gggtcggcgtcc---    602
                     ||.||     ||
Mouse            564 ggatc-----cccat    573
```

Suppl Fig S2

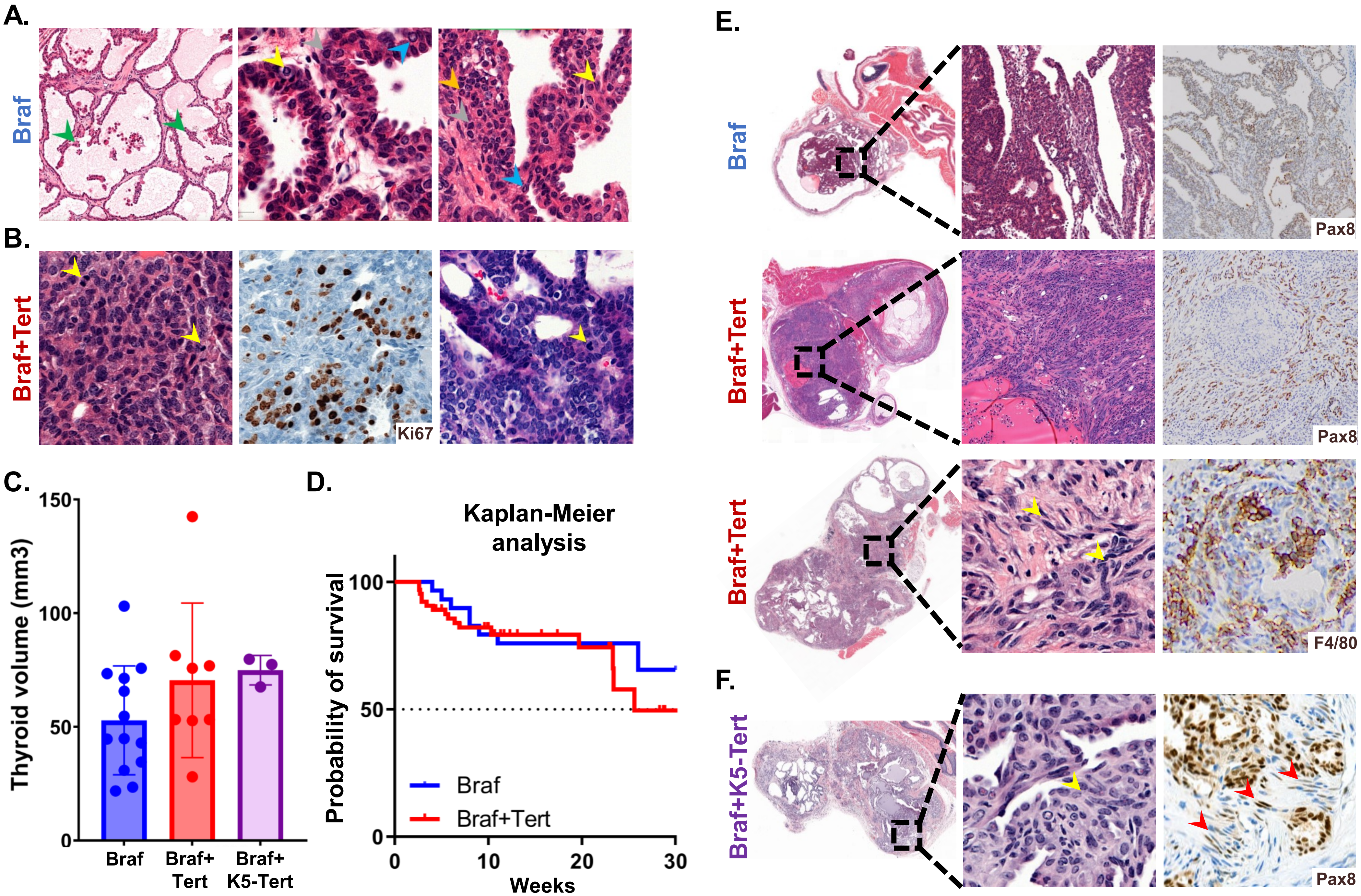
A.
Braf
B.
Braf+Tert
Ki67
C.
Thyroid volume (mm3)
150
100
50
0
Braf
Braf+
Tert
Braf+
K5-Tert
D.
Kaplan-Meier
analysis
Probability of survival
100
50
0
0
10
20
30
Weeks
Braf
Braf+Tert
E.
Braf
Pax8
Braf+Tert
Pax8
Braf+Tert
F4/80
F.
Braf+K5-Tert
Pax8

Suppl Fig S2, continued

G.

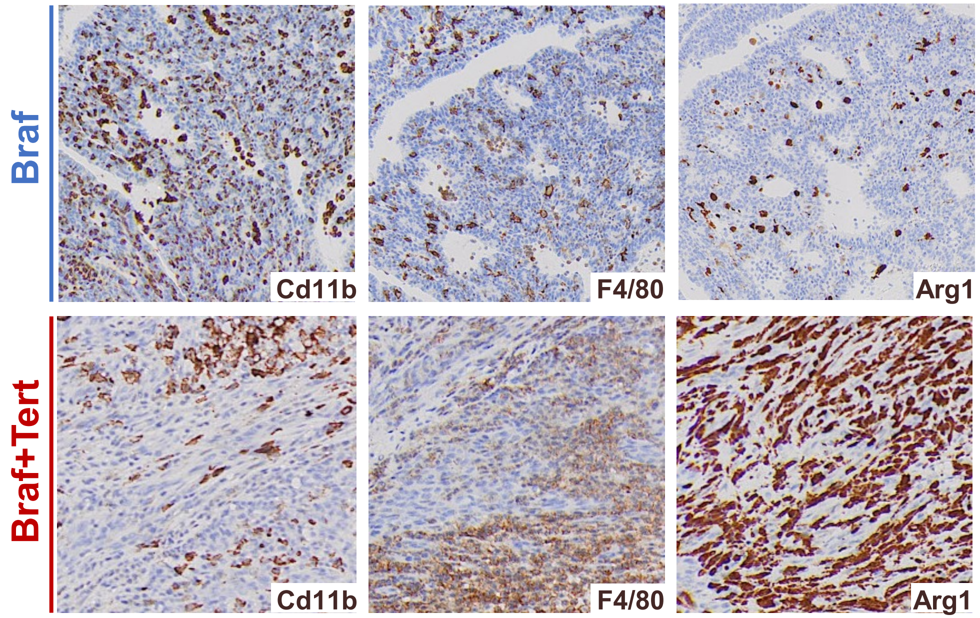


H.

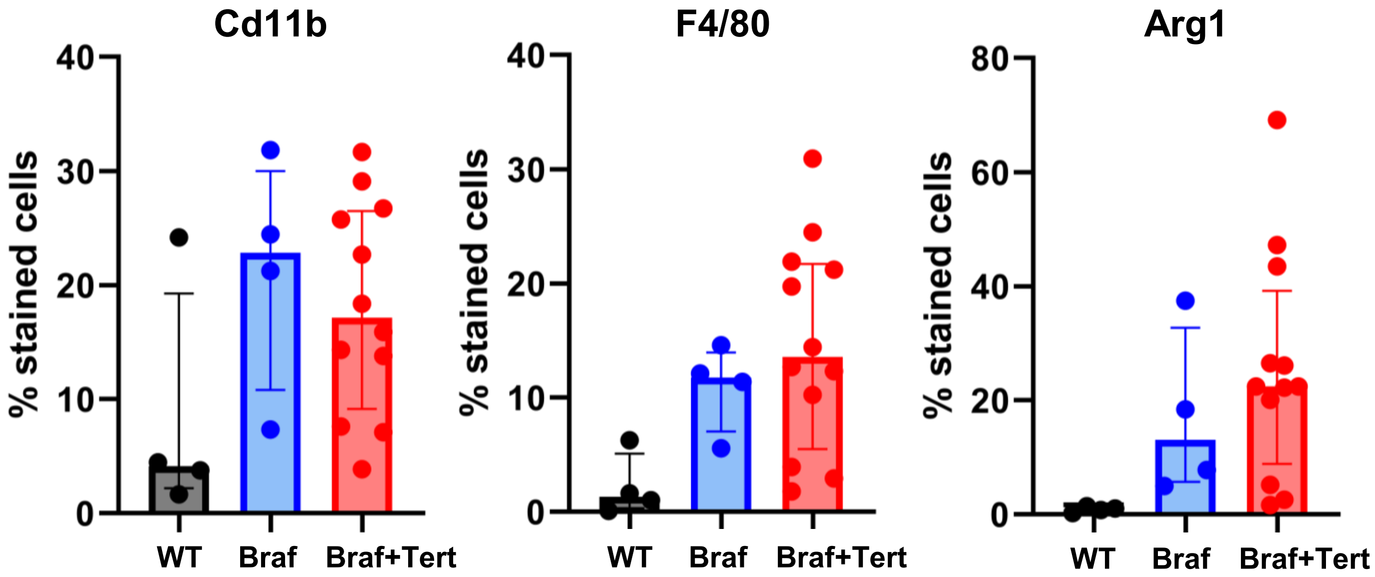

Suppl Fig S3

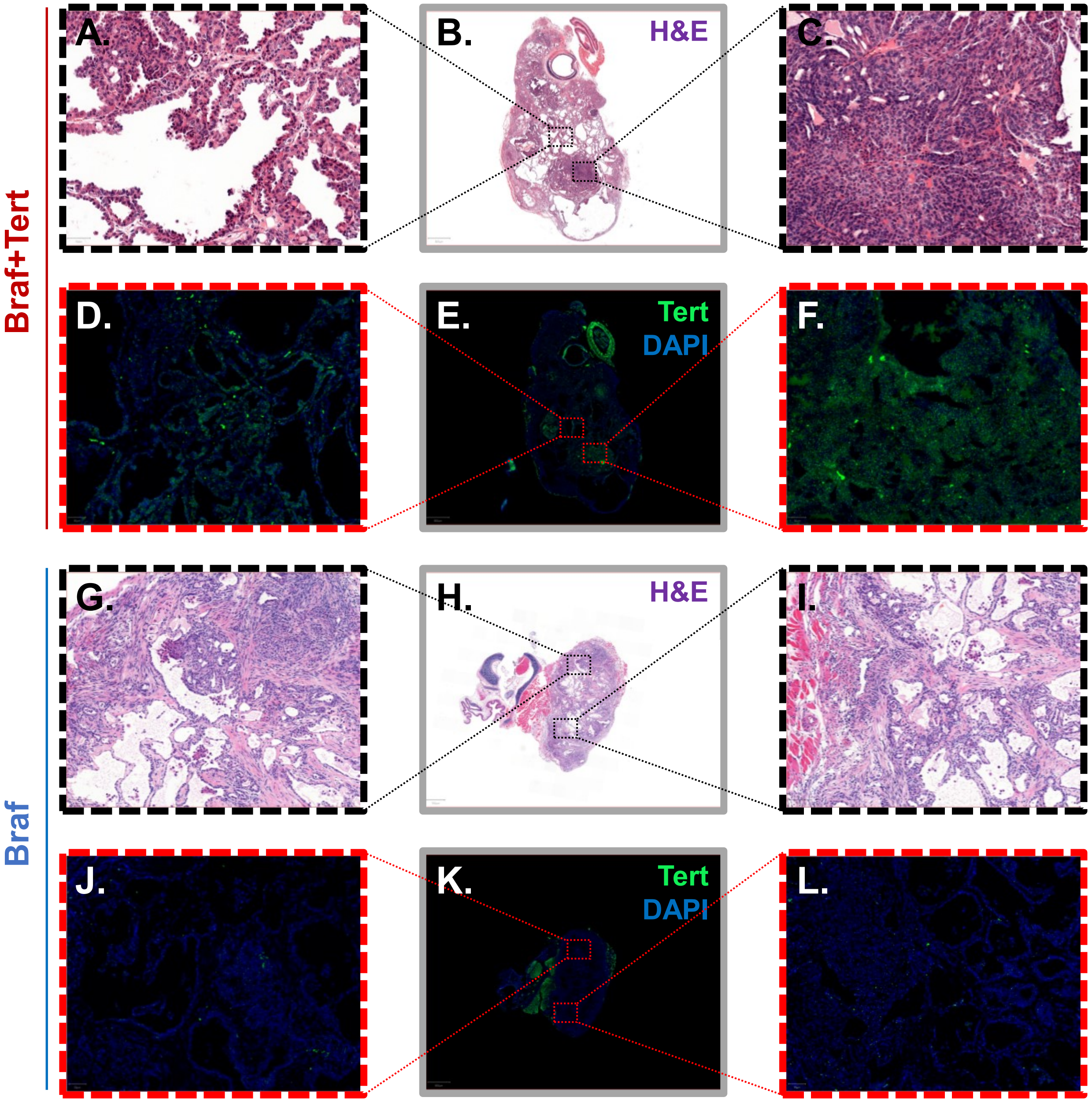

Suppl Fig S4

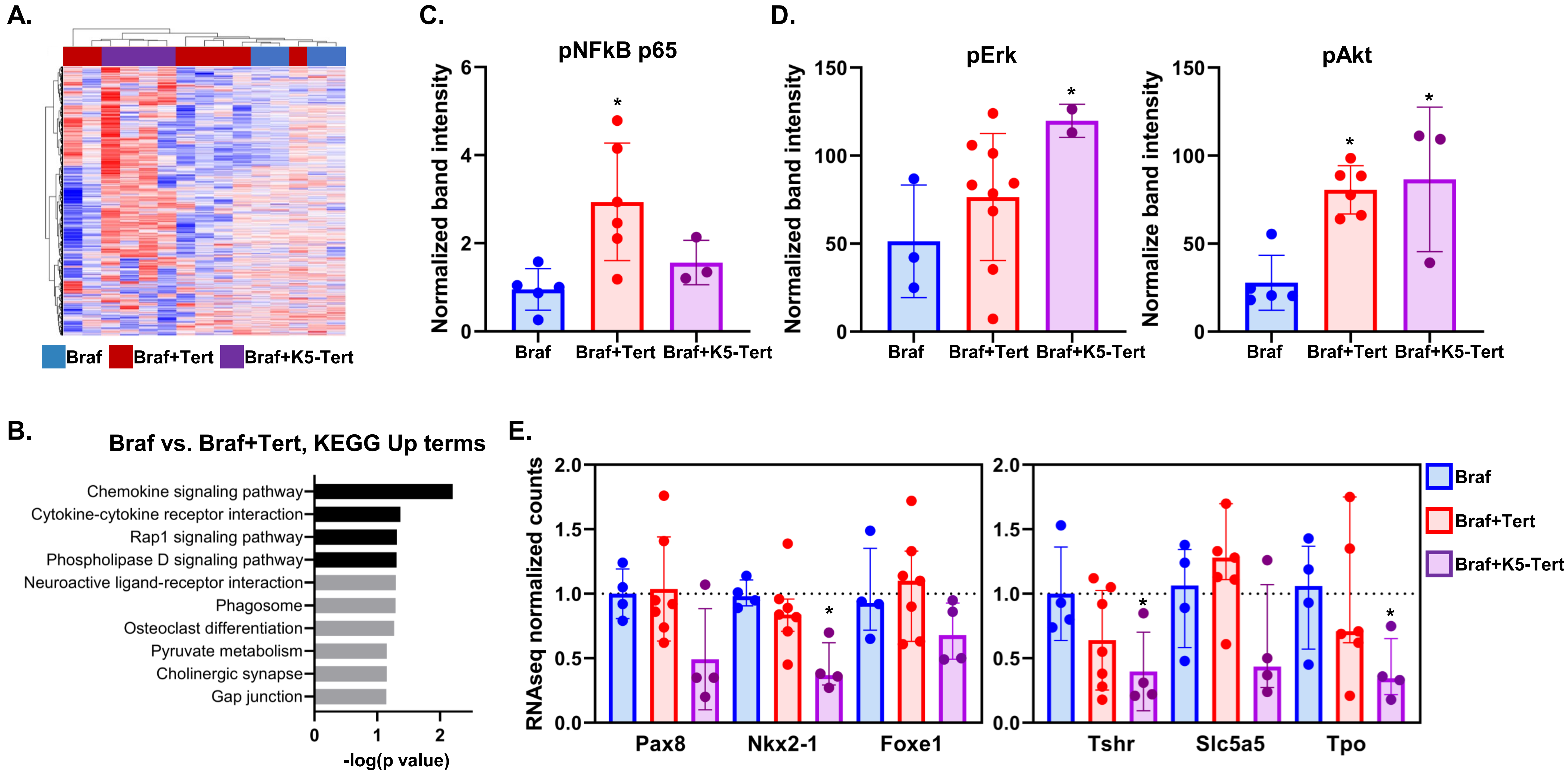
A.
Braf
Braf+Tert
Braf+K5-Tert
B.
Braf vs. Braf+Tert, KEGG Up terms
Chemokine signaling pathway
Cytokine-cytokine receptor interaction
Rap1 signaling pathway
Phospholipase D signaling pathway
Neuroactive ligand-receptor interaction
Phagosome
Osteoclast differentiation
Pyruvate metabolism
Cholinergic synapse
Gap junction
0
1
2
-log(p value)
C.
pNFkB p65
Normalized band intensity
0
2
4
6
*
Braf
Braf+Tert
Braf+K5-Tert
D.
pErk
Normalized band intensity
0
50
100
150
*
Braf
Braf+Tert
Braf+K5-Tert
pAkt
Normalize band intensity
0
50
100
150
*
*
Braf
Braf+Tert
Braf+K5-Tert
E.
RNAseq normalized counts
0.0
0.5
1.0
1.5
2.0
*
*
*
Pax8
Nkx2-1
Foxe1
Tshr
Slc5a5
Tpo
Braf
Braf+Tert
Braf+K5-Tert

Suppl Fig S5

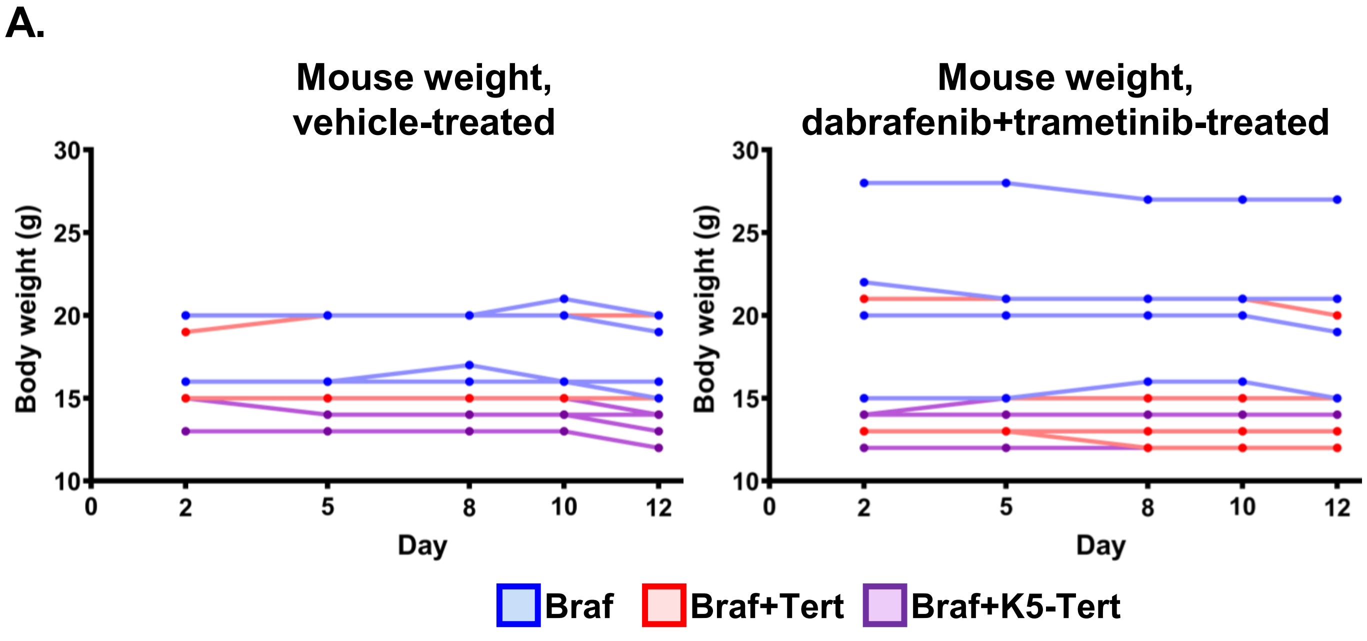
A.
Mouse weight, vehicle-treated
Mouse weight, dabrafenib+trametinib-treated
Body weight (g)
Day
Braf
Braf+Tert
Braf+K5-Tert

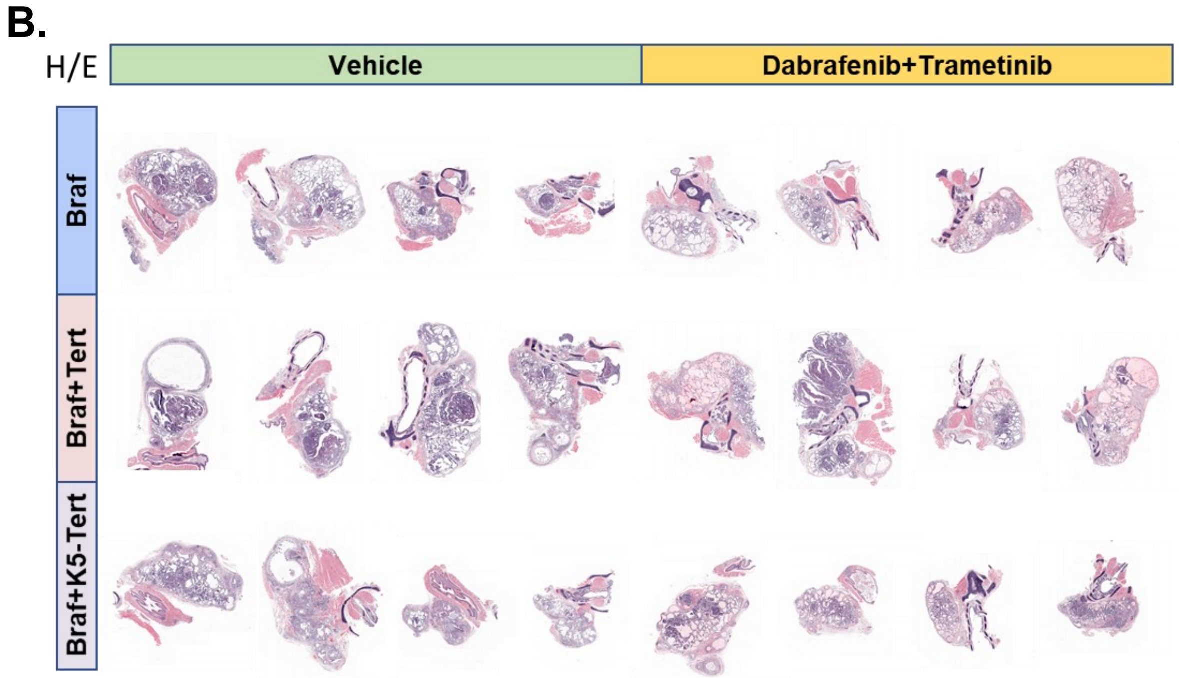
B.
H/E
Vehicle
Dabrafenib+Trametinib
Braf
Braf+Tert
Braf+K5-Tert

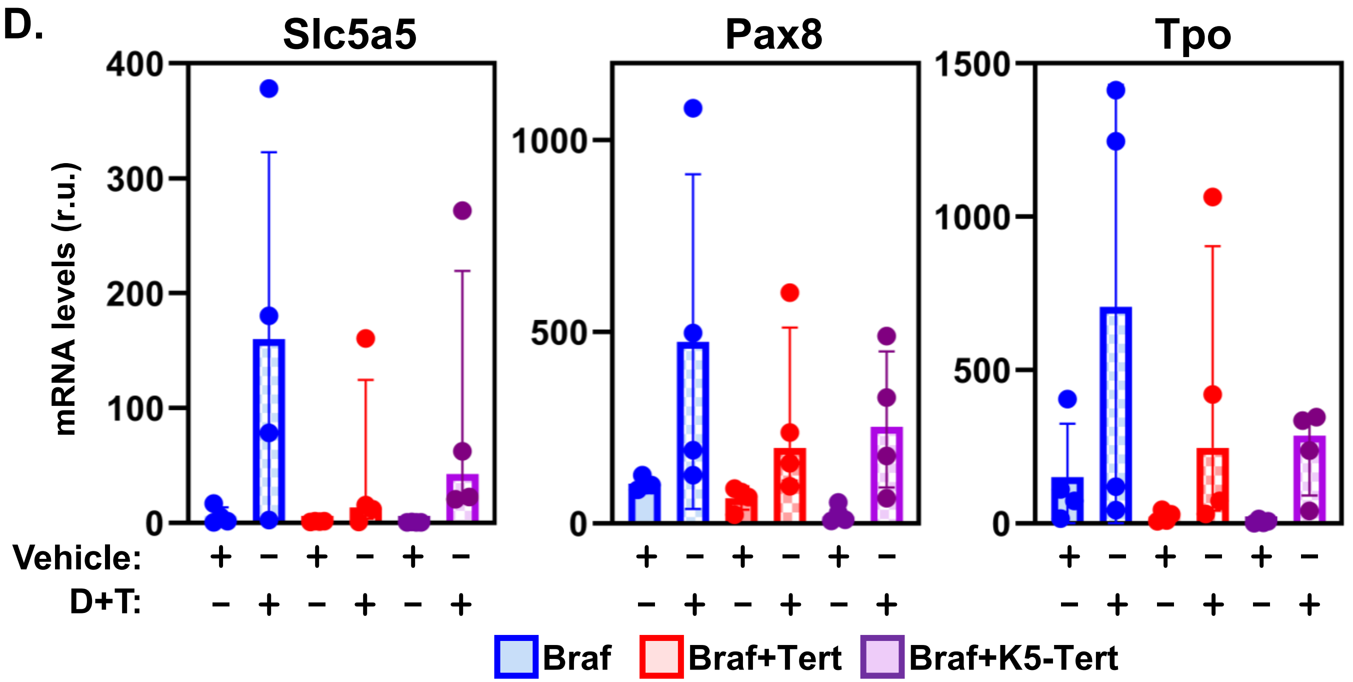
D.
Slc5a5
Pax8
Tpo
mRNA levels (r.u.)
Vehicle:
D+T:
Braf
Braf+Tert
Braf+K5-Tert

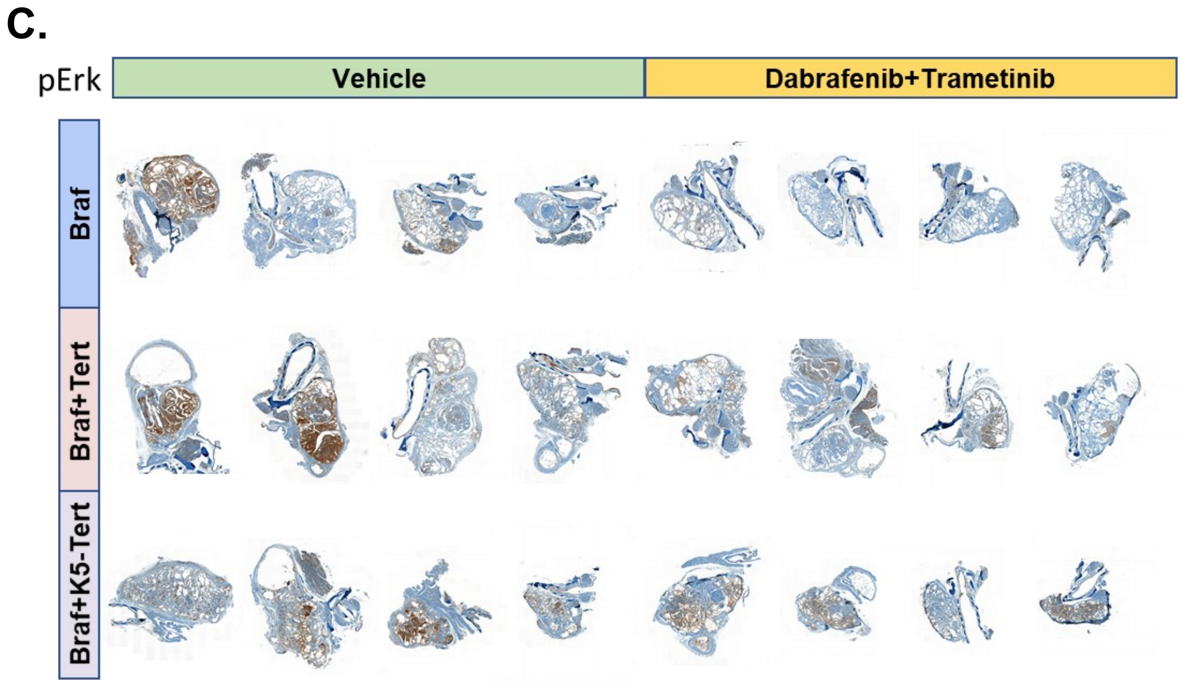
C.
pErk
Vehicle
Dabrafenib+Trametinib
Braf
Braf+Tert
Braf+K5-Tert
